# STILL-^13^C: Spatial tracing of isotopically labelled lipids with ^13^C reveals metabolic heterogeneity in intact tissues

**DOI:** 10.64898/2026.08.21.746222

**Authors:** Jacob X.M. Truong, Paul J. Trim, Jayden C. McKinnon, Kylie Taylor, Marten F. Snel, Shane R. Ellis, Johannes V. Swinnen, Lisa M. Butler

**Affiliations:** South Australian immunoGENomics Cancer Institute (SAiGENCI), Adelaide University Medical School, North Terrace, 5000, Adelaide, South Australia; Laboratory of Lipid Metabolism and Cancer, Leuven Institute for Single Cell Omics (LISCO) and Leuven Cancer Institute (LKI), KU Leuven 3000, Leuven, Belgium; South Australian Health and Medical Research Institute (SAHMRI), North Terrace, 5000, Adelaide, South Australia; Flinders Omics Facility, College of Medicine and Public Health, Flinders University, Sturt Road, Bedford Park, 5042, South Australia; Molecular Horizons and School of Science, Faculty of Health, Science and Medicine, University of Wollongong, Wollongong, 2500, NSW, Australia

## Abstract

Lipid metabolism is dynamically rewired across tissues in response to developmental, environmental and therapeutic cues. This adaptation drives treatment resistance in a range of human pathologies, but current lipidomic techniques fail to capture the underlying mechanisms, relying on steady-state measurements from homogenised samples that obscure spatial heterogeneity and pathway flux. Here we introduce spatial tracing of isotopically labelled lipids (STILL-^13^C), a workflow that uses stable isotope tracing and high-resolution mass spectrometry imaging (MSI) to map lipid metabolic flux directly in intact human tissues with unprecedented pathway coverage. STILL-^13^C overcomes longstanding limitations of bulk and MSI-based analyses by spatially resolving isotopologue labelling of simple and complex lipids, enabling simultaneous tracing of fatty acid synthesis, remodelling and multiple convergent pathways required for phospholipid assembly, while preserving tissue architecture and regional metabolic context. Applied to patient-derived prostate cancer explants cultured *ex vivo*, STILL-^13^C revealed heterogeneity in lipid pathway activity between neighbouring epithelial regions and spatially resolved responses to pathway inhibition. This work establishes a broadly applicable platform for investigating spatial heterogeneity in lipid metabolic flux and its perturbation in intact tissues.

## Main

Altered lipid metabolism has emerged as a central hallmark of numerous diseases, including cancer (1), neurodegeneration (2) and inflammatory conditions (3). As essential components of cellular membranes, energy storage and signalling pathways, lipids are intrinsically linked to a wide range of cellular processes. Consequently, targeting lipid metabolic pathways has become an increasingly promising strategy for improving patient outcomes (4). However, successful clinical translation will depend on technologies capable of capturing the full complexity of lipid metabolism *in situ*.

Lipid metabolism is highly dynamic, spatially organised and continuously rewired in response to environmental cues (5). These features are particularly pronounced in tumours, where cancer cells and stromal components engage in spatially heterogeneous and temporally evolving lipid exchange and signalling, generating regionally distinct metabolic states that cannot be resolved by bulk or static measurements of lipid profiles. Therefore, there is a critical need for approaches that can spatially resolve lipid dynamics in intact tissue contexts. Such tools would enable monitoring of patient response while simultaneously revealing metabolic adaptations that drive therapeutic resistance within the tumour and its microenvironment.

Stable isotope tracing with ^13^C partly addresses this need by measuring the incorporation of labelled substrates into newly synthesised metabolites. This approach has become a powerful tool for investigating lipid metabolism across diverse biological systems (6–8), but for tissues, has primarily been applied to bulk homogenates (9), which sacrifices spatial metabolic complexity. In contrast, matrix-assisted laser desorption/ionisation mass spectrometry imaging (MALDI-MSI) enables *in situ* analysis of metabolites across tissue sections, preserving tissue context, but has largely been restricted to static molecular profiling. MSI has been used to profile lipids in surgical specimens from multiple cancer types including endometrial (10), non-small cell lung (11), prostate (12–14), breast (15) and brain cancers (16). However, studies attempting *in situ* ^13^C tracing (17–24) remain limited and technically challenging, due to poor labelling efficiency, limited molecular coverage, isotopologue overlap and spatial labelling heterogeneity. Moreover, these approaches have rarely been applied in clinically relevant models or actual patient samples, leaving key aspects of human lipid metabolic plasticity unexplored.

To address these challenges, we developed STILL-^13^C (Spatial Tracing of Isotopically Labelled Lipids with ^13^C); a spatial tracer lipidomics workflow that measures lipid metabolic flux directly within intact human tissues. We apply this approach to prostate cancer patient-derived explants (PDEs), a clinically relevant model of primary disease (25) that enables controlled perturbation of metabolic pathways and inclusion of unlabelled controls for robust calculation of label enrichment. Prostate cancer displays a well-established dependency on lipid metabolism for growth, progression and survival, characterised by altered expression of lipid metabolic enzymes, such as those involved in *de novo* lipogenesis (26), fatty acid (FA) uptake/transport (27) and remodelling (28, 29), FA oxidation (30, 31), and phospholipid (PL) remodelling (14, 32, 33). Many of these are associated with aggressive disease and poor clinical outcomes (32, 33), making prostate cancer a compelling system for methodological validation.

In this work, we demonstrate the capacity of the STILL-^13^C workflow to spatially resolve lipid metabolic dynamics in fresh human prostate cancer tissues, establishing a generalisable framework for mapping multiple lipid metabolic pathways within complex tissue architectures. We further demonstrate clinical utility by monitoring treatment response and metabolic adaptation following inhibition of the key lipid metabolic enzyme fatty acid synthase (FASN). To our knowledge, this work represents the first application of spatial ^13^C-tracer lipidomics in clinical cancer specimens, addressing multiple longstanding technical challenges within a single integrated platform.

## Results

### Spatial Tracing of Isotopically Labelled Lipids with ^13^C (STILL-^13^C) workflow

To spatially map lipid dynamics in human prostate specimens, we performed MALDI-MSI on prostate PDEs from *N*=4 patients. Tissues were cultured in media supplemented with uniformly labelled ^13^C-glucose ([U-^13^C]-glucose) and treated with either vehicle control (DMSO) or the fatty acid synthase inhibitor TVB2640 (10 µM, FASNi) (Fig. 1). For each patient, matched explants cultured with unlabelled glucose (D-glucose) served as controls for endogenous unlabelled analytes. PDEs were processed according to a standard sample preparation workflow for MSI and analysed using a Bruker timsTOF fleX mass spectrometer, enabling high-resolution spatial profiling of labelled and unlabelled lipids.

**Fig. 1.**
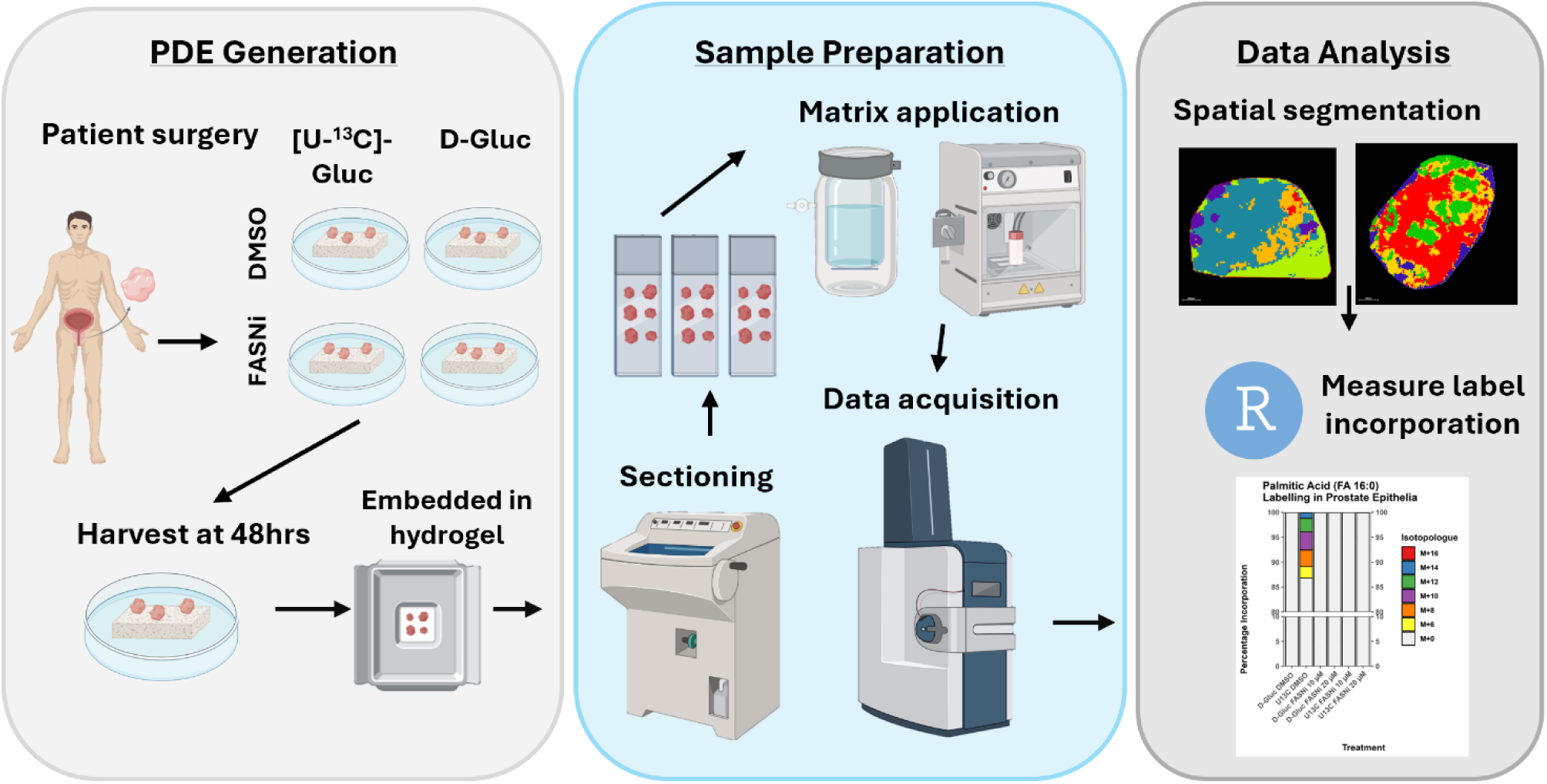
Experimental workflow for the spatial analysis of lipid dynamics in [U-^13^C]-glucose labelled prostate PDEs by MALDI-MSI. Surgical prostate cancer specimens were cultured as PDEs in media supplemented with uniformly ^13^C-labelled glucose ([U-^13^C]-Gluc) or unlabelled glucose (D-Gluc) and treated with vehicle control (DMSO) or a fatty acid synthase inhibitor (FASNi). Following culture, tissues were harvested, snap frozen, embedded, sectioned, and mounted onto glass slides. Sections were coated with a MALDI matrix and analysed by MALDI-MSI using a timsTOF fleX mass spectrometer. Data were subsequently processed to assess label incorporation into lipids using R-based analytical workflows.

### Tracing *de novo* fatty acid synthesis in prostate PDEs

Using the STILL-^13^C workflow in prostate PDEs we initially assessed ^13^C label incorporation into palmitate (FA 16:0), the 16-carbon end product of FASN activity, in its free form (Fig. 2). *De novo* fatty acid synthesis proceeds through iterative addition of two-carbon units derived from M+2 malonyl-CoA (where M+X indicates analyte M with X ^13^C labels) (Fig. 2A). Accordingly, [U-^13^C]-glucose-derived palmitate is measured as a series of even-numbered isotopologues. Odd-numbered isotopologues, attributable to natural ^13^C abundance (34), were excluded from downstream analyses. Spatial distributions of FA 16:0 M+2 and M+4 isotopologues differed substantially from higher-mass isotopologues (spearman *ρ* = 0.01 – 0.1), consistent with interference from endogenous unlabelled species in matched control tissues (Supplementary Fig. 1A, B), and previous observations in mouse brain (35). These were therefore excluded from enrichment analyses to avoid skewing of the isotopologue distribution. However, serial sections from two sets of patient-matched tissues imaged using a SolariX FT-ICR instrument confirmed that the M+4 isotopologue had endogenous interference that can be resolved using ultra high-resolution instruments (∼ 200k resolving power [FWHM] at *m/z* 259.24637 required for baseline resolution) (Supplementary Fig. 1B).

**Fig. 2.**
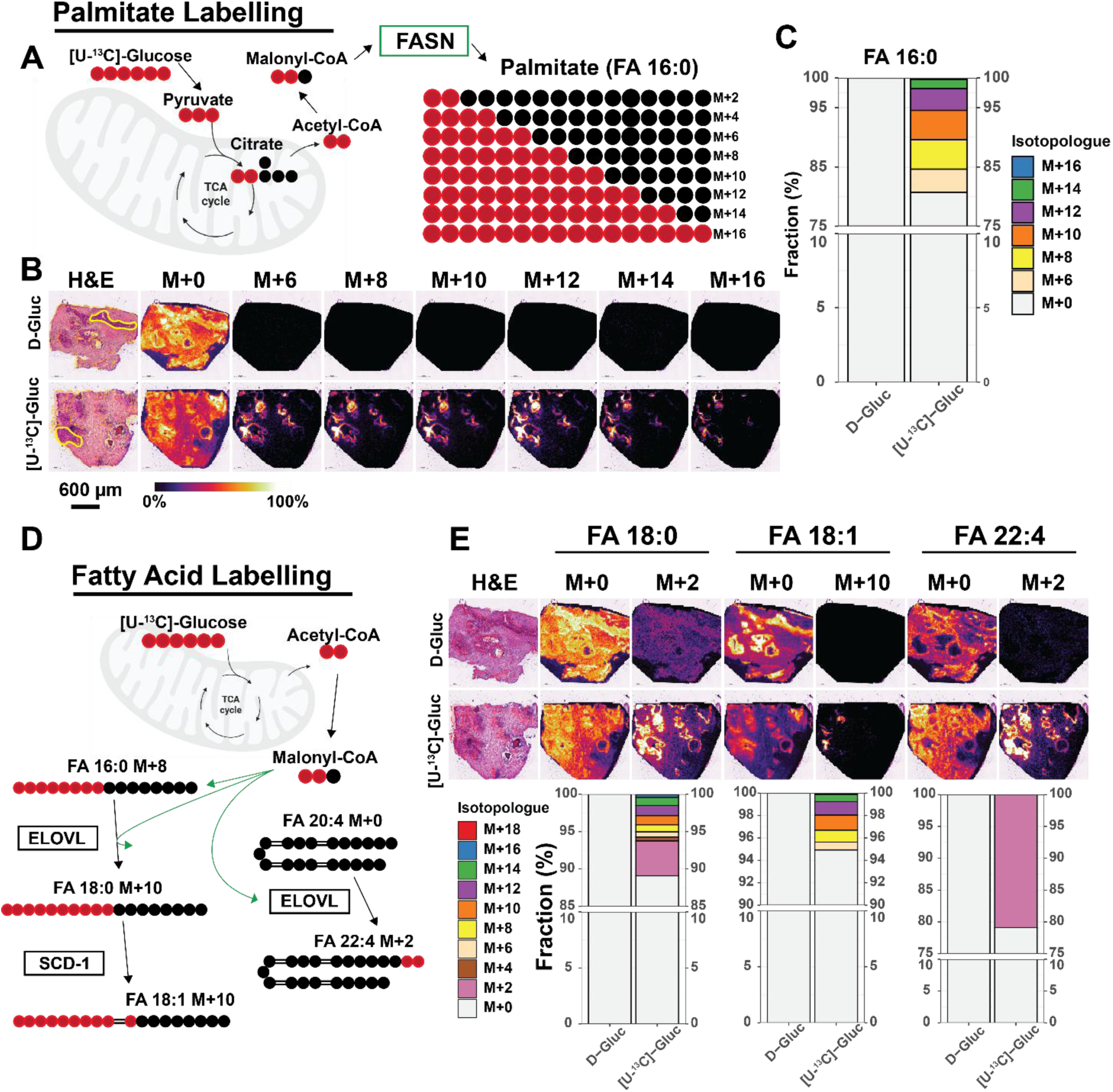
Human prostate PDEs incorporate [U-^13^C]-glucose into fatty acids in the glandular epithelium. **A)** Schematic of FA 16:0 (palmitate) synthesis by FASN, showing stepwise incorporation of two ^13^C atoms from lipogenic acetyl-CoA. Created with BioRender.com. **B)** Ion distribution images of M+0 and even-numbered isotopologues of FA 16:0 in matched prostate PDE tissues cultured in media supplemented with D-glucose (top row) or [U-^13^C]-glucose (bottom row). Corresponding H&E-stained section indicates glandular epithelium and stromal regions. **C)** Isotopologue enrichment plot showing fractional contributions to the free FA 16:0 pool in the glandular epithelium tissue. **D)** Schematic of metabolic pathways utilising labelled acetyl-CoA derived from [U-^13^C]-glucose for the synthesis and elongation of FAs. Created with BioRender.com. **E)** Representative isotopologue images and corresponding enrichment plots for FA 18:0, FA 18:1 and FA 22:4 in matched tissues cultured in media supplemented with D-glucose (top row) or [U-^13^C]-glucose (bottom row). All ion images are intensity scaled within a single *m/z* channel. Enrichment plots are derived from natural ^13^C abundance corrected data from *N*=4 patients after removing isotopologues with endogenous overlap.

Spatial clustering of MSI data across the four patients consistently identified glandular epithelium and stroma for downstream enrichment analysis, each exhibiting distinct labelling profiles (Supplementary Fig. 2A). To measure average FA 16:0 enrichment within individual epithelia glands, the average peak area of each isotopologue (excluding M+2 and M+4), was extracted from regions defined on the corresponding H&E image (Fig. 2B). This analysis was performed across all patients, and isotopologue enrichment was calculated following correction for natural ^13^C abundance of the even isotopologues (Fig. 2C). On average, 19.3% +/- 5.2% of total palmitate across the cohort was measured as ^13^C-labelled, with the isotopologue distribution centred around M+10 at 5.0% (SD = 1.5%). This suggests ∼60% of the acetyl-CoA pool was labelled after 48h of culture, in line with previous ^13^C tracing studies (34). No labelled isotopologues (M+6 to M+16) were detected in matched D-glucose control tissues (Fig. 2B), indicating negligible interference from endogenous unlabelled species. In ^13^C-glucose samples, *de novo* synthesis of longer chain fatty acids was evident in epithelial regions, with stearate (FA 18:0), FA 18:1 and FA 22:4 measured in their free form showing average enrichments across the cohort of 11.1% ± 1.8%, 5.3% ± 2.7% and 20.7% ± 5.7% respectively (Fig. 2E). Notably, whereas the endogenous unlabelled pool of FA 22:4 (M+0) was higher in stromal regions relative to glandular epithelium, the only labelled adrenate isotopologue detected (M+2) was restricted to epithelial regions, similar to the other labelled FAs (Fig. 2E). After isotopic correction, the highest fractional enrichment for FA 18:0 and FA 22:4 was measured for the M+2 isotopologues (4.8% ± 0.7% and 19.5% ± 6.3% respectively). These labelling patterns suggest that FA 18:0 and FA 22:4 were predominantly generated via elongation of unlabelled FA 16:0 and FA 20:4 respectively, using labelled acetyl-CoA (M+2) as a substrate (Fig. 2D). However, despite FA 18:0 M+2 being the most enriched, no signal was detected for the M+2 isotopologue of FA 18:1 in any sample. A complete table with corrected isotopologue fractions for each fatty acid isotopologue series from all tissues analysed is included in Supplementary File 2.

### *In situ* mapping of complex lipid assembly and metabolism

Free FAs are essential building blocks of more complex phospholipids. Therefore, to visualise the dynamics of phospholipid synthesis and measure FA incorporation, we investigated isotopic labelling of multiple phospholipid and lysophospholipid classes including PI, LPI, PA, PC, LPC, SM, PE, LPE, PG and PS. For several lipids, specific isotopologue *m/z* channels exhibited interference from endogenous unlabelled species present in matched tissues cultured in D-glucose medium and were therefore excluded from enrichment analyses. Multiple phospholipids showed distinct spatial distributions for the labelled species compared to the unlabelled (M+0). Examples of those measured with high abundance in PDE tissues are depicted in Fig. 3A. PI lipid signals were strongly enriched in glandular epithelium relative to the surrounding stroma, whereas PE 36:1 and PC 34:1 were detected in both compartments and SM d34:1 intensity was higher in stromal regions. Despite these differences in M+0 signal distributions, newly labelled isotopologue signals (shown for M+14) were consistently enriched in glandular epithelium for all these lipids (Fig 3A). This mirrored the spatial patterns observed for *de novo* synthesised FAs in Fig. 2E, indicating higher relative rates of local *de novo* synthesis in glandular regions compared to stroma. Isotope labelling patterns of these lipids (Fig. 3B and Supplementary Fig. 2C) revealed both even-numbered isotopologues resulting from *de novo* FA synthesis and odd-numbered isotopologues arising from labelling of glycerol-3-phosphate (G3P) in the phospholipid backbone (8, 9, 36). After isotope correction, the M+3 isotopologue was dominant across all measured phospholipids (Fig. 3B and Supplementary Fig. 2C). Among sphingomyelins, only SM d34:1 showed evidence of ^13^C FA labelling, potentially reflecting a lower turnover in stroma and therefore an insufficient labelling duration for pathways involved in sphingolipid metabolism. Importantly, we reproduced these spatial labelling patterns across multiple instruments (Thermo Orbitrap Elite and Bruker SolariX FT-ICR) and laboratories (Supplementary Fig. 2D, E) validating the reliability and generalisability of STILL-^13^C. Fractional enrichments for all samples acquired on each MSI platform are provided in Supplementary File 3.

**Fig. 3.**
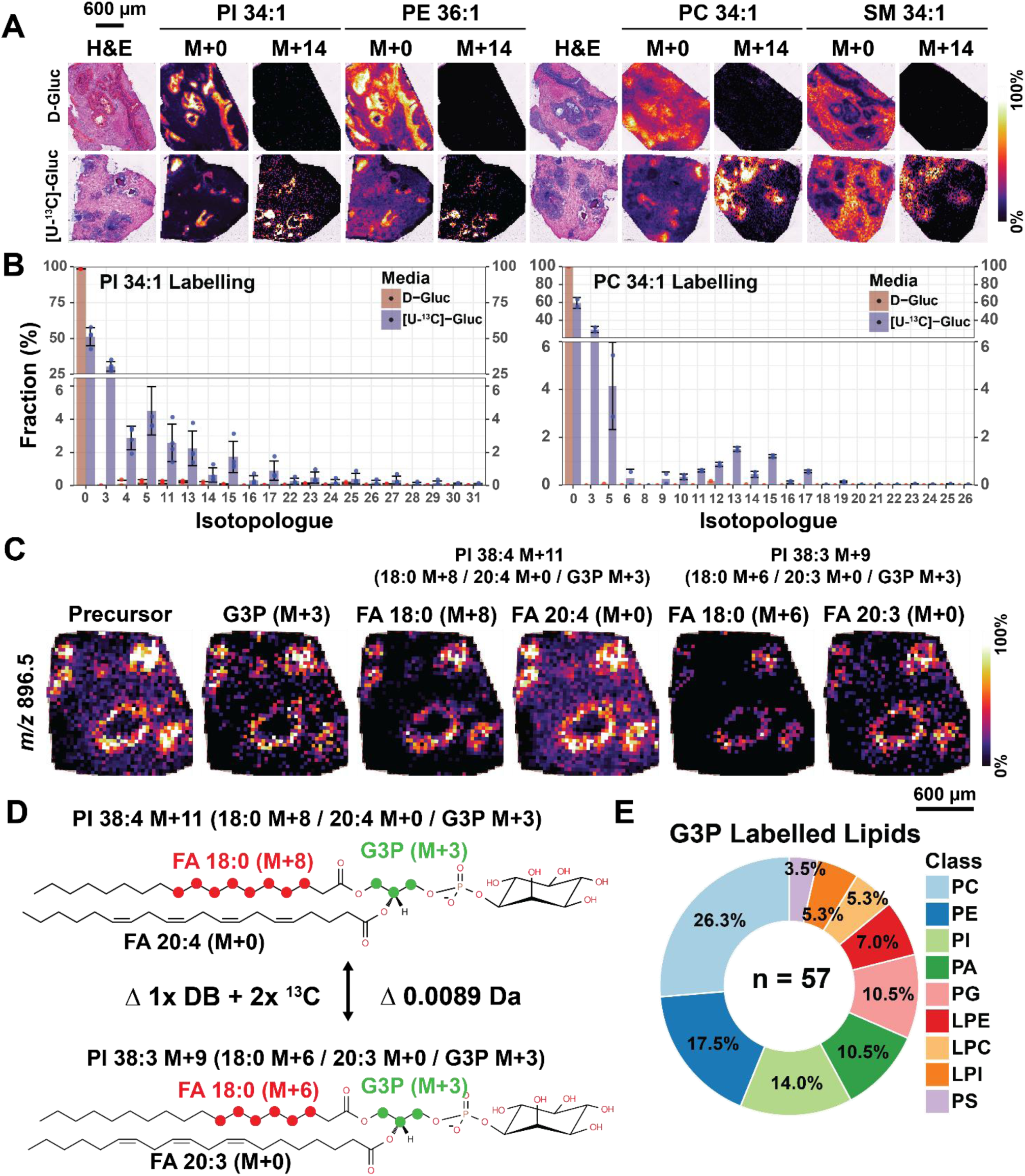
Prostate epithelia incorporate *de novo* synthesised FAs into phospholipids and utilise glucose-derived ^13^C to synthesise G3P. **A)** Ion distribution images of the unlabelled and M+14 isotopologues of PI 34:1 and PE 36:1 detected in negative ion mode and PC 34:1 and SM d34:1 measured in positive ion mode in matched tissues cultured in D-glucose or [U-^13^C]-glucose supplemented media. **B)** Isotopologue enrichment plots for PI 34:1 and PC 34:1 showing the labelling patterns across the cohort in both medium conditions. Isotopologue fractions represent the mean fraction (%) ± SD of *N*=4 patients in negative ion mode and *N*=3 patients in positive ion mode. **C)** Fragment ion distribution images from MS/MS imaging acquisitions of PI 38:4 M+11 and PI 38:3 M+9 showing labelling of G3P and FA chains. **D)** Chemical structures showing the FA species and labelling states of annotated lipids from MALDI-MS/MS imaging of precursor ion *m/z* 896.5868 and the mass offset introduced through ^13^C labelling. Double bond location and FA *sn* position were not experimentally determined. **E)** Pie chart showing the number of lipids detected as the M+3 isotopologue (or M+6 for PG lipids) and their percentage contribution per subclass. All ion images are intensity scaled to within a single *m/z* channel and enrichment plots are derived from natural ^13^C abundance corrected data after removing isotopologues with endogenous overlap. DB = double bond.

To demonstrate spatial lipid annotation beyond the sum composition (aggregate carbons and double bonds), we performed MALDI-MS/MS imaging for PI 38:4 as the M+11 isotopologue in two patient samples, alongside a D-glucose treated control tissue. The precursor ion (*m/z* 896.5868) and the G3P M+3 fragment (*m/z* 156.006) localised to glandular epithelium (Fig. 3C and Supplementary Fig. 3A) based on post-acquisition H&E staining (Supplementary Fig. 3A). The dominant FA fragment pair associated with PI 38:4 M+11 comprised FA 18:0 M+8 and FA 20:4 M+0, together with G3P M+3, all co-localising with the precursor ion signal (Fig. 3C). The isolated precursor at *m/z* 896.5868 overlapped with a PI 38:3 M+9 isotopologue of nearly identical mass (*m/z* 896.5957, Δ0.0089 Da) which differs from PI 38:4 M+11 by one double bond in the unlabelled C20 FA and two ¹³C labels in FA 18:0 (Fig. 3D, labelled moieties highlighted). Such near isobaric overlaps are an inherent challenge in ^13^C labelling experiments and while these can be resolved in the lower fatty acid mass range (*m/z* 200-350) (Supplementary Fig. 4A), they can only be resolved in the phospholipid mass range (*m/z* 600-900) using high mass-resolution MS platforms such as an Orbitrap or FT-ICR (Supplementary Fig. 4B, C), or by using orthogonal separation techniques such as ion mobility spectrometry. To demonstrate this, we performed MSI on serial sections using trapped ion mobility spectrometry (TIMS) and showed comparable separation of these isobars (Supplementary Fig 4D). MS/MS data did not support labelling of FA 20:4. However, multiple labelled states of FA 18:0 (M+4 to M+12) and FA 20:3 (M+2) co-localized with the precursor (Supplementary Fig. 3A). Overall, MS/MS imaging identified seven distinct labelled lipid species combining unlabelled FA 20:3/20:4 with variably labelled FA 18:0 and G3P (Supplementary File 4), all of which were spatially resolved to glands, indicating both *de novo* synthesised FAs and those derived through other pathways (fatty acid uptake) are used for phospholipid synthesis. This pattern of multiple labelling states was consistent with on-tissue MS/MS profiling data of different PI 38:4 and PI 34:1 isotopologues (Supplementary Fig. 5, 6). While label incorporation into fatty acyl chains was only observed in the most abundant phospholipids, labelling of the G3P backbone was widespread across a broad range of lipid classes. In particular, M+3 or M+6 for PG isotopologues (6) (which contain two glycerol moieties) as confirmed by MS/MS, were detected in numerous lipid subclasses, most prominently in PCs, followed by PEs and PIs (Fig. 3E).

### STILL-^13^C resolves spatial heterogeneity in FA synthesis and elongation in histologically similar glandular epithelia

The differing spatial patterns between FA isotopologues (Fig. 2) and labelled FA fragment ions in glandular epithelium (Fig. 3C) suggest metabolic heterogeneity within this tissue compartment. To assess whether additional *de novo* synthesised FAs were labelled but undetected in free form, we performed all-ion fragmentation (AIF) imaging (negative mode, *m/z* 100–1000, CID) without precursor selection, as described previously (21). Tissues (*N* = 2) cultured in [U-¹³C]-glucose were analysed alongside matched D-glucose controls (Supplementary Fig. 7). Indeed, AIF revealed labelled fatty acids that were missed by direct MSI (Fig. 2). This enabled spatial tracing of stepwise elongation for C20–C24 saturated FAs (SFAs) from unlabelled FA 18:0 (Fig. 4A) and desaturation of FA 18:0 to FA 18:1 (M+10), followed by elongation to C20–C24 monounsaturated FAs (MUFAs; Fig. 4B). Notably, the distribution of newly elongated products differed: labelled C20-C24 SFAs were broadly distributed throughout most of the epithelia, whereas elongated MUFAs localised to discrete glands. Even within the same tissue, AIF data revealed marked differences between histologically similar glands (Fig. 4C). In one representative gland (“gland 1”), lighter isotopologues of FA 16:0 and FA 18:0 predominated, whereas heavier isotopologues were enriched in an adjacent “gland 2” (Fig. 4C). Correspondingly, isotopologue distributions shifted from M+6 to M+10 for FA 16:0 and from M+10 to M+14 for FA 18:0 (Fig. 4D). Together, these data demonstrate substantial metabolic heterogeneity across morphologically similar prostate glands and highlight the ability of MSI to spatially resolve differences in lipid metabolic pathways.

**Fig. 4.**
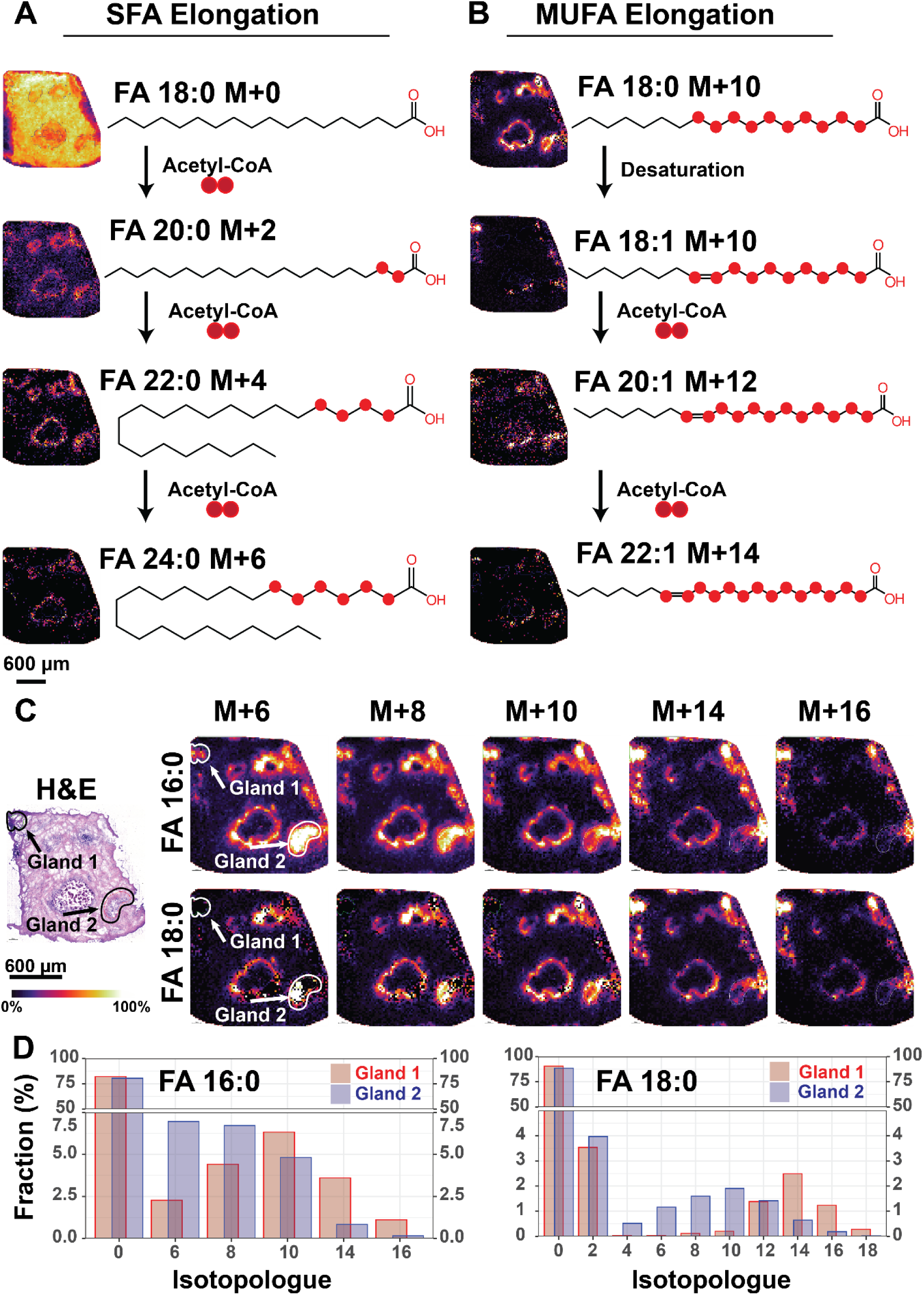
MALDI-MSI resolves spatial heterogeneity in FA 16:0 synthesis and elongation of C18-C24 SFAs and MUFAs in histologically comparable glandular epithelia. **A)** and **B)** Ion distribution images and annotated structures showing step-wise elongation via sequential addition of two ^13^C units from acetyl-CoA for **A)** C18-C24 SFAs and **B)** C18-C22 MUFAs. **C)** Post-acquisition H&E staining and ion distribution images of the total labelled FA 16:0 and FA 18:0 pool liberated through AIF imaging. **D)** Isotopologue distributions for FA 16:0 (left) and FA 18:0 (right) overlaid for regions corresponding to glands annotated in **C)**. Ion distribution images are normalised within each *m/z* channel. Enrichment plots are derived from natural ^13^C abundance corrected data after removing isotopologues with endogenous overlap.

### STILL-^13^C resolves metabolic responses to FASN inhibition

To assess how spatial ^13^C-lipidomics can reveal treatment response and lipid metabolic plasticity upon lipid pathway perturbation in patient samples, we applied the STILL-^13^C workflow to spatially map metabolic responses to FASN inhibition in prostate PDEs. Tissues were cultured for 48hrs with 10µM TVB2640, a FASN inhibitor under clinical Phase I investigation for castration-resistant prostate cancer (CRPC) (clinical trial number: NCT05743621) (37). FASN inhibition markedly suppressed incorporation of [U-^13^C]-glucose-derived carbon into newly synthesised fatty acids and phospholipids across epithelial regions, while preserving labelling patterns consistent with elongation of pre-existing FAs and glycerol backbone synthesis. Isotopologue fractions were significantly decreased by over 98% in the glandular epithelium of TVB2640-treated tissues compared to DMSO controls (Wilcoxon Rank Sum *p* < 0.05) for all isotopologues of FA 16:0 (Fig. 5A), FA 18:1 and FA 18:0, except for FA 18:0 M+2 (Supplementary Fig. 7A & Supplementary File 5) and the only measured isotopologue of FA 22:4 (M+2) (Fig. 5B). Retention of the M+2 isotopologues of FA 18:0 and FA 22:4 suggests continued elongation of unlabelled FAs via elongation activity independent of FASN. Phospholipid labelling was similarly reduced following FASN inhibition. Fig. 5C shows isotopologue distributions for PI 34:1 across DMSO- and TVB2640-treated tissues cultured in [U-¹³C]-glucose, with representative ion images for M+0, M+3 and M+17. Fractional enrichment decreased significantly (*p* < 0.05) for all isotopologues except M+3, M+4, M+5, M+27 and M+28. Notably, the persistence of M+3 and M+5 isotopologues across multiple phospholipid classes (Supplementary Fig. 7B; Supplementary File 5) indicates that G3P labelling from glucose was unaffected by FASN inhibition, as expected. Consistent with this interpretation, on-tissue targeted MS/MS analysis of TVB2640-treated tissues detected no labelled phospholipid precursor ions or labelled FA fragments (Supplementary Fig. 5, 6), confirming selective inhibition of phospholipid synthesis pathways dependent on newly synthesised FAs.

**Fig. 5.**
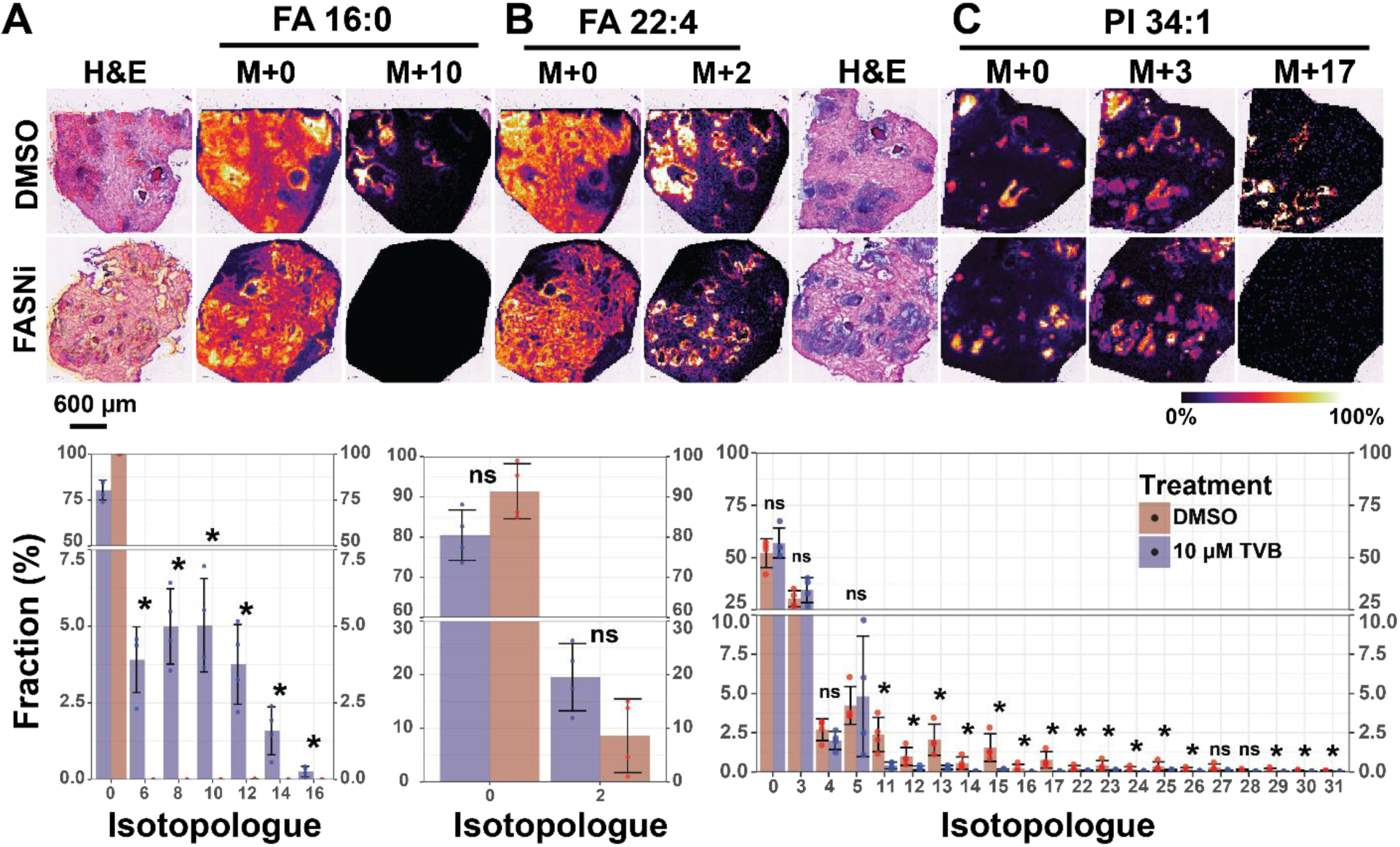
Pharmacological inhibition of FASN reduces incorporation of [U-^13^C]-glucose into *de novo* synthesised fatty acids and phospholipids. A-C) Post MALDI H&E staining and ion distribution images of **A)** FA 16:0 M+10, **B)** FA 22:4 M+2 and **C)** PI 34:1 M+3 and M+17, with corresponding isotopologue distributions comparing mean fractions of isotopologues between tissues cultured in [U-^13^C]-glucose and treated with DMSO (red) or 10 µM TVB2640 (blue). Ion distribution images are normalised within each *m/z* channel. Enrichment plots are derived from natural ^13^C abundance corrected data after removing isotopologues with endogenous overlap.

## Discussion

In this study, we present STILL-^13^C, a spatial ^13^C-tracer lipidomics method capable of mapping lipid metabolic flux and pathway perturbation directly in intact human tissues. Using this approach, we show that lipid metabolic activity in human prostate cancer tissue is spatially compartmentalised, pathway-selective and only partly predicted by histology or steady-state lipid abundance. Our workflow demonstrates the capability of STILL-^13^C for direct visualisation of metabolic pathway activity within intact tissue architecture, linking pathway dynamics to histological features and revealing heterogeneity that would be obscured by bulk analyses.

Applied to prostate patient-derived explants (PDEs), which retain tissue architecture and cellular heterogeneity (25), this workflow resolves multiple interconnected lipid metabolic pathways. Using [U-^13^C]-glucose, we spatially map *de novo* fatty acid synthesis, desaturation, elongation – all processes implicated in prostate cancer progression (28, 29, 38–40) and therapeutic targeting, as well as phospholipid production. These analyses demonstrate that lipid metabolic flux is far from uniform; newly synthesised or modified FAs and phospholipids were concentrated in epithelial regions while adjacent stromal regions showed minimal ^13^C labelling, consistent with prior evidence of higher glycerol-3-phosphate dehydrogenase 1 (GPD1), FASN and ELOVL isoform expression in prostate epithelia tissue compared to stroma (28, 29, 41, 42). Moreover, this pattern did not simply mirror the distribution of the corresponding unlabelled lipids. For example, FA 22:4 and SM d34:1 showed discordant distributions between unlabelled and labelled pools, demonstrating that static lipid abundance can misrepresent the spatial sites of active lipid synthesis or remodelling. We further observed significant heterogeneity even across histologically similar individual epithelial glands, underscoring the importance of spatially resolved measurements for understanding metabolic organisation and plasticity. These findings suggest that neighbouring histological regions or cell types can have distinct metabolic states, each with potentially different therapeutic vulnerabilities.

Beyond fatty acid synthesis, STILL-^13^C enables simultaneous interrogation of glycerophospholipid assembly by capturing labelling in both fatty acyl chains and the glycerophosphate backbone. The predominance of M+3 isotopologues across phospholipid classes indicates preferential routing of glucose-derived carbon into glycerol-3-phosphate via glycolysis, whereas heavier isotopologues arising from combined fatty acid and backbone labelling are spatially restricted. The proportions of M+3 labelled phospholipids revealed highest backbone metabolism in PC, PE and PI lipids, consistent with their enrichment in malignant tissue (12, 43–45). Combined backbone and FA labelling in phospholipids coupled with undetectable FA labelling in sphingolipids beyond SM d34:1 also indicates preferential routing of newly synthesised FAs toward phospholipid assembly in proliferative prostate epithelia, although extended labelling times and alternative tracers such as [U- ^13^C_3_]-serine (9) may be required to fully resolve/capture sphingolipid flux. Nevertheless, our data support that complex lipid production is modular and spatially heterogeneous in intact tissues, and that retention of spatial information is critical to disentangle these parallel metabolic processes.

A key strength of the workflow is the integration of complementary MSI modalities. All-ion fragmentation (AIF) and targeted MS/MS imaging expand molecular coverage by recovering fatty acid fragments from complex lipids and enabling direct interrogation of isotopic labelling within lipid species. This reveals spatially distinct labelling patterns, including regions with inverse patterns of FA labelling (e.g. differential labelling of FA 16:0 and FA 18:0 between different glands), indicative of local variation in lipogenic flux that would be averaged out in bulk analyses. Resolving elongation and desaturation products further enables inference of pathway activity and substrate flow.

Untargeted MSI and AIF revealed labelling of FA 16:0, FA 18:0 and FA 18:1, but not FA 16:1, consistent with preferential utilisation of newly synthesised FA 16:0 for elongation over desaturation, despite the reported upregulation of both pathways in primary disease (28, 29, 39). Although double-bond position was not directly resolved, this pattern is consistent with labelled FA 18:1 being predominantly oleic acid, the *n*-9 isomer synthesised by ELOVL6 elongation of FA 16:0 (M+X) to FA 18:0 (M+[X+2]) (46) and subsequent desaturation by SCD-1 (28). Elongation of unlabelled FA 16:1 into the FA 18:1 *n*-7 isomer, *cis*-vaccenic acid (M+2), which also plays an oncogenic role in prostate cancer (28) by ELOVL5 is possible, however FA 18:1 M+2 was not detected in any sample.

Recovery of labelled SFAs and MUFAs by AIF also supports increased carbon routing through elongation pathways as products are already incorporated in phospholipids at 48hrs. These examples illustrate how STILL-^13^C can provide functional metabolic insight using widely accessible instrumentation.

Importantly, this method supports interrogation of metabolic perturbations in patient samples, expanding its utility beyond basic discovery and into clinical translation. Using FASN inhibition as an example, we demonstrate selective suppression of *de novo* fatty acid synthesis while preserving labelling in pathways independent of FASN activity. Persistence of M+2 fatty acid isotopologues (FA 18:0, FA 22:4) and M+3 phospholipid isotopologues in TVB2640-treated tissues further confirms that fatty acid and phospholipid synthesis in PDEs is both modular and pathway-convergent. While sustained fatty acid synthesis/remodelling using FA 16:0 is suppressed, there is continued elongation and phospholipid assembly using pre-existing fatty acids potentially supplied through elevated exogenous fatty acid uptake (47). These conserved spatial labelling patterns indicate heterogeneity in adaptive lipid re-routing pathways across complex tissues that would be obscured by measuring static lipid abundances in bulk and which could reveal candidate combinatorial therapeutic targets.

Combined with the ability of MSI to map drug distribution within tissues, which has previously been demonstrated in prostate PDEs (48), STILL-^13^C provides a means of correlating drug distributions/uptake with metabolic pathway adaptation in clinically relevant systems. This integrated approach could ultimately help identify metabolic predictors of drug response or resistance in a clinically-relevant model setting.

Although demonstrated here using ^13^C-glucose, the framework is readily extendable to alternative ^13^C tracers such as acetate or glutamine, enabling interrogation of multiple metabolic inputs into lipid biosynthesis that shift under different environmental conditions (49). Incorporating additional substrates, perturbations or time-resolved measurements (e.g. shorter timepoints to capture initial flux kinetics) and computational flux modelling may further expand throughput and pathway resolution.

Technically, the method addresses key challenges in the field. In contrast to recent applications of tracer lipidomics, which have mapped metabolic flux and response to lipid pathway perturbation in bulk tissue homogenates (9), the implementation of MSI technology here resolved these dynamics in a complex tissue and in doing so revealed spatial heterogeneity in these processes. The STILL- ^13^C method also addresses limitations with recently reported spatial tracer methods (18–20, 50). For example, isobaric overlap with unlabelled lipids are accounted for by culturing patient-matched tissues in regular medium. Tracing from central carbon metabolism into simple and complex lipids is possible due to high label incorporation in the PDE model while targeted MS/MS imaging improves molecular annotation, confidence in isotopologue assignment, and can spatially resolve heterogenous labelling patterns. Furthermore, complementary platform comparisons highlight the benefit of high-mass resolution MS or orthogonal separation techniques in resolving isobaric overlaps, although the increased acquisition time and data file size of ultra-high resolution systems introduce limitations for scalability. Despite the advances that the STILL-^13^C method provides to the field, there are some limitations to our method. Steady-state labelling is unlikely to be achieved during the 48-hour culture period, making pathway dynamics difficult to model as previously demonstrated (9), however prostate PDEs show viability up to 96 hours (25) and have been treated for up to 72 hours (51). Limits in sensitivity restricted MSI and MS/MS imaging experiments to 20 µm and 40 µm spatial resolutions respectively, although post-ionisation MALDI-2 technology may recover losses in sensitivity at higher spatial resolutions, allowing for finer detailed metabolic mapping (52, 53).

Overall, this work establishes STILL-^13^C as a broadly applicable platform for profiling metabolic flux within intact patient-derived samples, revealing spatial pathway organisation that is invisible to both bulk tracer lipidomics and static MSI, providing a powerful foundation for advancing metabolism-focused precision medicine. This capability opens new opportunities to investigate metabolic heterogeneity, cell state plasticity and therapeutic adaptation across cancer and other diseases where metabolism is spatially organised.

## Methods

### Reagents

Red phosphorous, N-(3-Dimethylaminopropyl)-N′-ethylcarbodiimide hydrochloride (NEDC), 2,5-dihyroxybenzoic acid (DHB), Norharmane, sodium formate, DPX mountant, Anti-mycotic, Dimethyl sulfoxide (DMSO), Polyvinylpyrrolidone, (Hydroxypropyl)methyl cellulose and [U-^13^C]-glucose were purchased from Sigma-Aldrich (MO, USA). Methanol, ethanol, xylene, hydrochloric acid (36%), acetonitrile, 2-propanol (IPA), acetone and LC-MS grade water were purchased from Chem Supply (SA, Australia). Filtered Lillie-Mayer’s Haematoxylin and 1% Alcoholic Eosin were purchased from Australian Biostain (VIC, Australia). ESI-L Tune Mix and Bluing buffer were purchased from Agilent (CA, USA). M4 culture medium containing 10% (v/v) dialysed serum and unlabelled glucose solution (D-glucose) were purchased from Thermo Fisher Scientific (MA, USA). TVB2640 was purchased from Sapphire Bioscience (NSW, Australia).

### Tissue collection

Radical prostatectomy (RP) samples for explanting were collected from *N* = 4 consenting patients undergoing surgery at St Andrew’s Hospital (Adelaide, Australia) with permission from the Human Research Ethics Committees of St Andrew’s Hospital (Approval #80) and the University of Adelaide (Approval #H-2023-067). After surgery, the removed prostate was placed in a sterile container on ice and a single core from the left and right lobe of the prostate was removed by a pathologist using either a 6 mm or 8 mm biopsy punch at the site of cancer according to the biopsy report. The tissue cores were placed into tubes and transported on ice for processing.

### Prostate patient-derived explants

Explants were prepared as previously described by Centenera et al. 2018 (25), under a project ethics approval from the University of Adelaide (#H-2012-016). Briefly, tissue cores from surgery were dissected into 4 x 1 mm^3^ pieces and placed onto gelatin Spongostan sponges (McFarlane Medical, VIC, Australia) suspended in 500 µL of glucose free M4 culture medium containing 10% (v/v) dialysed serum and 1% (v/v) antimycotic in a sterile plastic 24-well culture plate. For each patient, explants were cultured for 48hrs in media supplemented with either 1.11 mM D-Glucose or [U-^13^C]-Glucose and treated with either dimethyl sulfoxide (DMSO) as a control, or 10 µM TVB-2640 (FASN inhibitor). The timepoint of 48 hours was chosen to allow substantial label incorporation while maintaining tissue viability. At 24hrs, media from all samples was stripped and replaced with fresh media. All explants were cultured at 37 °C in a 5% CO_2_ incubator (Bio Strategy, NSW, Australia).

After 48hrs, tissues were placed in plastic 10×10×5 mm cryomolds (Thermo Fisher Scientific, MA, USA) and embedded in 500 µL of hydrogel (7.5 g (w/v) hydroxypropyl methylcellulose, 2.5 g (w/v) polyvinylpyrrolidone) (54), immediately frozen on dry ice and stored at -80 °C until processing.

### Tissue sectioning

All sectioning was performed using a Leica CM 1950 (Leica Biosystems, Nussloch, Germany). PDE samples were removed from -80°C storage and placed in the cryostat chamber for a minimum of 30 minutes prior to sectioning at -20°C. Once tissues were warmed to -20°C they were sectioned at 10 µm onto indium-tin oxide coated slides (Bruker Daltonics). Serial sections were collected on consecutive slides for small molecule metabolite imaging, positive ion mode lipidomics and negative ion lipidomics. Each slide contained a single section from each matched set of PDEs, this allowed all treatment groups to be compared on a single slide for each analysis type reducing potential batch effects caused by sample preparation and data acquisition. An additional serial section was taken onto a Superfrost Plus microscope slide for H&E staining (Thermo Fisher, MA, USA).

### Matrix application for fatty acid imaging (NEDC)

Slides used for negative ion mode small molecule metabolite imaging were spray coated with NEDC matrix according to the methods published by Kasarla *et al* 2025 (55). Briefly, slides were thawed to room temperature in a vacuum desiccator for 30 minutes prior to opening to prevent condensation. A freshly prepared solution of 7 mg/mL NEDC matrix in 70:25:5 MeOH:MeCN:H2O (v/v/v) was prepared and briefly sonicated. A total of 21 layers of matrix was applied using a SunCollect (SunChrom, Germany) sprayer, for all layers the *z* position of the sprayer was maintained at 35mm. The first 3 layers were applied at a flow rate of 5 µL/min followed by 3 layers at 10 µL/min, 3 layers at 15 µL/min and the final 11 layers at 20 µL/min. With an *x* and *y* axis speed of 620 and 850 mm/s respectively. A fixed pressure of nitrogen gas was supplied at 2.5 bar.

### Matrix application for lipid imaging (DHB and Norharmane)

2,5-DHB or norharmane were sublimated onto slides for positive and negative ion mode lipid imaging respectively using our previously published method (56). Briefly, 7.5 mg of DHB or norharmane matrix was dissolved in 200 µL acetone (DHB) or 200 µL of methanol (norharmane) by sonication before being deposited onto a 90 mm diameter foil disk forming a uniform layer of matrix crystals. For norharmane, the foil disk was pre-warmed to 60 °C for 10 minutes and then placed in the base of a glass sublimation chamber (ACE Glass Incorporated, USA). For each application, up to 2 slides were mounted at the base of the cold finger and sealed. The system was evacuated by connecting it to a Freeze Drier system (Martin Christ, Germany) which was maintained at 0.12 mBar and -86°C for the ice condenser. The slides were then cooled by filling the cold finger with ice water. After allowing the temperature to equilibrate across the slide for 3 minutes, the matrix was heated using a heating mantal (Chiltern Scientific, Whangarei, New Zealand) to a maximum of 120°C over the course of 10 minutes. After sublimation was complete the heating mantel was switched off, followed by warming the cold finger to room temperature. The pressure in the chamber was slowly increased to atmospheric pressure, reducing the possibility of condensation on the sample. Matrix was then recrystalised by sealing the slides in a glass chamber containing 300 µL of 5% methanol at 60°C for 90 seconds.

### Free fatty acid imaging, timsTOF fleX

Small molecule imaging was performed in negative ion mode on a Bruker timsTOF fleX mass spectrometer (Bruker Daltonics, Bremen, Germany). Data was acquired between *m/z* 35-650 using a 20 µm pixel and step size. The Bruker SmartBeam^TM^ laser system was operated at 10 kHz with 80% laser fluency and 500 shots/pixel. To optimise the transfer of small molecules (*m/z* < 500), the MALDI plate offset was set to 50 V, with the funnel 1, funnel 2 and multipole RF voltages set to 125 V_pp_, 200 V_pp_ and 200 V_pp_ respectively. The collision RF voltage was set to 450 V_pp_. TOF detection was tuned for this mass range by setting the transfer time and pre-pulse storage to 55 µs and 5 µs respectively. Mass calibration was performed by direct infusion of a 1:1 mixture of ESI-L Tune Mix and 2 mM sodium formate in 50% IPA while operating the instrument in ESI mode. The instrument syringe pump was operated at 3 µL/min, and mass calibration was performed using the high precision calibration (HPC) mode. The calibration settings were accepted when an accuracy score of at least 98% was achieved with a standard deviation of < 1ppm.

### Lipid imaging, timsTOF fleX

Lipid imaging in positive and negative ion mode was performed on the timsTOF fleX with an adjusted method for the detection of phospholipids. For lipids in positive and negative ion mode, data was acquired between *m/z* 300-1000 and *m/z* 200-100 respectively using the same laser spot size of 20 µm but with 250 shots/pixel. Transfer of ions in the lipid mass range was optimised by setting the RF voltages in funnel 1, funnel 2, the multipole and the collision cell to 350 V_pp_, 350 V_pp_, 300 V_pp_ and 1800 V_pp_ respectively for both methods. The transfer time and pre-pulse storage was set to 85 µs and 10 µs respectively for both methods.

On-tissue MALDI-MS/MS data was acquired between *m/z* 100-1000 in negative ion mode. Acquisitions were performed using the sweeping acquisition tool in timsControl, allowing the instrument to partition the total laser shots between multiple tune profiles. The deflection delta and RF voltages for funnel 1, funnel 2 and the multipole were switched to 300 V_pp_, 350 V_pp_ and 400 V_pp_ respectively. A total of 500 shots were used per acquisition using a 40×40 µm laser spot size at 65% laser fluency. For 30% of the 500 shots, the collision RF, transfer time and collision energy were held at 1800 V_pp_, 80 µs and 10 eV to acquire precursor ion data. These parameters were adjusted to 800 V_pp_, 40 µs and 55 eV for the remaining 70% of the sweeping acquisition to acquire fragmentation data. For all MS/MS acquisitions, a pre-pulse storage of 10 µs was used and 2000 laser shots were summed over four acquisitions (500 shots/acquisition) using a ± 1.5 Da quadrupole isolation width. The exact same instrument settings were used for MS/MS imaging experiments.

All ion fragmentation (AIF) was performed with the same parameters as indicated above for MS/MS acquisitions with the following adjustments. The number of laser shots/pixel was decreased to 250 and the laser spot size was dropped to 20 µm. The collision energy and transfer time was maintained for the entire acquisition at 55 eV and 45 µs respectively. The instrument was operated in MS mode to ensure no precursor selection and online *m/z* calibration was performed using FA 18:1 M+0 (*m/z* 281.2486) as a reference mass with a minimum peak intensity of 5000 per spectra.

### Lipid imaging, Orbitrap

Norharmane matrix was dissolved in 2:1 chloroform:MeOH (v:v) to a concentration of 7 mg/mL. Samples were spray coated with an automated sprayer (TM-Sprayer, HTX Technologies, LLC, Chapel Hill, NC, USA). 15 layers of matrix were applied at a flow rate of 0.12 μL/min, nozzle temperature of 30°C, a velocity of 1200 mm/min and nitrogen gas pressure was maintained at 10 psi. The matrix density across the sample was 0.0035 mg/mm^2^. Data were acquired on an Orbitrap Elite mass spectrometer (Thermo Fisher Scientific GmbH, Bremen, Germany) equipped with an intermediate-pressure MALDI source (Spectroglyph LLC, WA, USA). A frequency-tripled Nd:YLF laser (Explorer One, Spectra Physics, Mountain View, CA) operating at 349 nm and 500 Hz was used for MALDI. The laser was run at a diode current of 1.9 A, and pulse energy was fine-tuned using an external attenuator (PowerXP, Altechna, Vilnius, Lithuania) to give a pulse energy entering the ion source of ∼ 1 µJ. All MSI data were acquired in negative ion mode at 240,000 Full Width at Half Maximum (FWHM) resolving power (at *m/z* 400) with a 300 ms injection time, resulting in a scan rate of approximately 0.97 scans per second. Data was collected from a mass range of *m/z* 350-1500. Raw data acquired using the Orbitrap Elite were recalibrated using Xcalibur RecalOffline software (ver. 4.1.50, Thermo Fisher Scientific). Theoretical values for [PS 36:1 - H]- (*m/z* 788.5447) and [PI 38:4 - H]- (*m/z* 885.5499) were used to recalibrate the data as these *m/z* values were of high relative abundance within majority of scans. Recalibration was conducted per scan with a mass tolerance of ±15 ppm. Final mass accuracy after recalibration is typically better than 2 ppm.

### Free fatty acid and lipid imaging in negative ion mode, 7T SolariX 2xR

Sample preparation was as described above in “matrix application lipid imaging”. Slides were imaged on a Bruker 7T SolariX 2xR (Bruker Daltonics, Bremen, Germany) using a Smart beam II laser operated at 2 kHz with 80% and 250 shots/pixel with laser focus set to minimum. Magnitude was set to 4M and the *m/z* acquisition range to *m/z* 43 – 1000. The capillary exit, funnel 1 and detector plate voltages were set to 200 V, 150 V and 220 V respectively. The RF voltage in the octupole and collision cell were set to 350 V_pp_ and 1800 V_pp_ respectively. Mass calibration was performed by MALDI using red phosphorus and the HPC mode. The final mass accuracy was ∼ 2.22 ppm (averaged over measurements for PA 36:1, PE 36:1, PE 38:4, PS 36:1 and PI 38:4) and the resulting mass resolution was 265,000 FWHM at *m/z* 400.

### H&E staining

Tissues imaged by MALDI-MSI were stained by first clearing the matrix with six washes in an ethanol gradient. Slides were submerged for 15 seconds in three solutions of 10%, 50% and 75% ethanol, followed by a reverse gradient in the same solutions before hydrating in a gentle stream of tap water for 1 minute. Tissues were then rinsed for 20 seconds in MilliQ water before staining in filtered Lillie-Mayer’s Haematoxylin for 45 seconds before rinsing in a gentle stream of tap water for 1 minute. The slides were blued by dipping in fresh 0.3% acid alcohol (HCl:Ethanol:H_2_O 0.3:70:30) two times followed by an additional three dips in 100% ethanol. Tissues were stained in 1%

Alcoholic Eosin for 15 seconds. Excess eosin was cleared by submerging slides in three solutions of 100% ethanol for three minutes each before submerging in two solutions of xylene for three minutes each. All slides were mounted with DPex, coverslipped and scanned on a Hamamtsu Nanozoomer 2 Digital Slide Scanner (Shizuoka, Japan). Additional H&E staining was performed as previously described (57) with the addition of a bluing step using Bluing buffer for 5 seconds after haematoxylin exposure or was performed at institutes where validation experiments were conducted using a modified protocol for prostate tissue (58).

### Data Processing in SCiLS Lab

All MSI datasets regardless of instrument were imported and processed in SCiLS Lab MVS 2025c (Bruker Daltonics, Bremen, Germany). Recalibrated Orbitrap data was imported as imzML files while data acquired on Bruker platforms was imported as native Bruker datatypes. Datasets were TIC normalised, and interval processing widths of ± 10 ppm, ± 7 ppm and ± 5 ppm was used with the Peak Area processing mode for timsTOF fleX, Orbitrap Elite and FT-ICR data respectively.

Three curated lists containing the theoretical *m/z* values of the target analytes and all their isotopologues were used as feature lists. The free fatty acid target list contained species with even numbered carbons from C14 – C24 and double bonds from 0 - 6. The negative ion mode lipids target list contained curated species from the PI, PE, PS, PG, LPE, LPI and LPA classes with sum carbon compositions consistent with containing FA chains from the small molecule target list. The feature list used for positive ion mode contained LPC and PC lipids with the same FA combinations. All target lists can be found in (Supplementary File 6). Regions of interest (ROIs) were defined in SCiLS Lab by comparing a spatial segmentation to the H&E image. Segmentation was performed using the bisecting *k*-means algorithm on TIC normalised data with the corresponding feature list for that imaging acquisition. The Manhattan distance metric was used with weak denoising. All tissues, regardless of the medium in which they were cultured in or the treatment, were segmented with the complete feature list containing the monoisotopic peaks with all theoretical isotopologues. The average peak area for each defined ROI was exported from SCiLS Lab for further processing and isotope correction.

### Isotope Correction

Correction for natural abundance of ^13^C isotopes was performed using IsoCor V2.2.3 (59). A metabolite database was generated for each acquisition using the chemical formulas of all targets in the curated feature lists. For the correction of timsTOF fleX QTOF, Thermo Orbitrap Elite and Bruker SolariX FT-ICR data, the resolution was set 40000 (constant), 240,000 (at *m/z* 400, Orbitrap) and 265,000 (at *m/z* 400, FT-ICR) respectively. Correction of the natural abundance of the tracer element (^13^C) was selected, and the tracer purity was set to 1. Original and corrected peak areas from all datasets are included in Supplementary File 7.

### R

Isotopologue enrichment calculations and data visualisation was performed using the *tidyverse* (V 2.0.0) packages in *R* (V 4.5.2) and RStudio (2026.01.0). Isotopologues series for each feature were first screened for interference from endogenous unlabelled analytes by assessment of ion images and mass spectra from unlabelled control tissues in SCiLS Lab. Once each isotopologue series had been screened and filtered to remove those with interference, fractional enrichments were calculated on the remaining isotopologues for each tissue and each lipid by calculating the proportion of the isotopologue peak area to the sum peak area of all isotopologues. An average enrichment per FA/lipid was calculated for all patients by averaging the data from each replicate tissue imaged. Statistically significant differences between fractional enrichments for each isotopologue between DMSO and TVB2640 treated samples were calculated using Wilcoxon Rank Sum test. Significance was reported as as *, *p* < 0.05; **, *p* < 0.01; ***, *p* < 0.001; *p* < 0.0001. All isotopologue enrichment plots represent the mean ± SD of *N*=4 patients in the cohort.

## Data Availability

Vendor neutral imzML files for all associated MSI data will be uploaded to public repositories.

## Acknowledgements

This work was supported by The Australian Research Council (DP230103210 to LMB and JVS) and the National Health and Medical Research Council of Australia (ID 2038959 to LMB and JVS), FWO grant G032525N (to JVS and LB), FWO-SBO S001623N (to JVS) and the Leuven Opening the Future Campaign. JXMT is supported by Leuven Future Fund LISCO-BIOMED. The South Australian immunoGENomics Cancer Institute (SAiGENCI) received grant funding from the Australian Government. S.R.E. acknowledges financial support the Australian Research Council (FT190100082). J.C.M and K. T are supported by Australian Government Research Training Program Scholarships

The authors acknowledge the team of the prostate cancer research group (Adelaide University, Adelaide, South Australia) and clinical staff of South Terrace Urology, Clinpath pathology, SA Pathology and St Andrew’s Hospital for coordinating the collection and culturing of patient tissues and the patients themselves for the donation of their tissue. We also wish to thank Dr Melvin Gay (Bruker) for technical support, advice and instrument setup of the 7T SolariX 2xR and to acknowledge the Australian National Phenome Centre for access to this instrument.

## Author contributions

JXMT and PJT contributed equally to the development of the method including data acquisition, data analysis, figure generation and manuscript preparation. JM, KT and SRE performed part of the mass spectrometry acquisition. LMB contributed to project conception, development, provision/coordination of samples, manuscript writing and supervision. JVS contributed to project conception, manuscript writing and supervision. MFS and PJT contributed to theoretical modelling of high-resolution data. PJT, MFS, SRE and JVS contributed to maintenance of instrumentation.

## Ethics declarations

The authors declare no competing interests.

