## Supplementary File 1 for "STILL-^13^C: Spatial tracing of isotopically labelled lipids with ^13^C reveals metabolic heterogeneity in intact tissues"

First authors

* Corresponding authors:

\# Current address: The Kinghorn Cancer Centre, The Garvan Institute of Medical Research, Sydney, 2010, NSW, Australia

**Supplementary File 1**

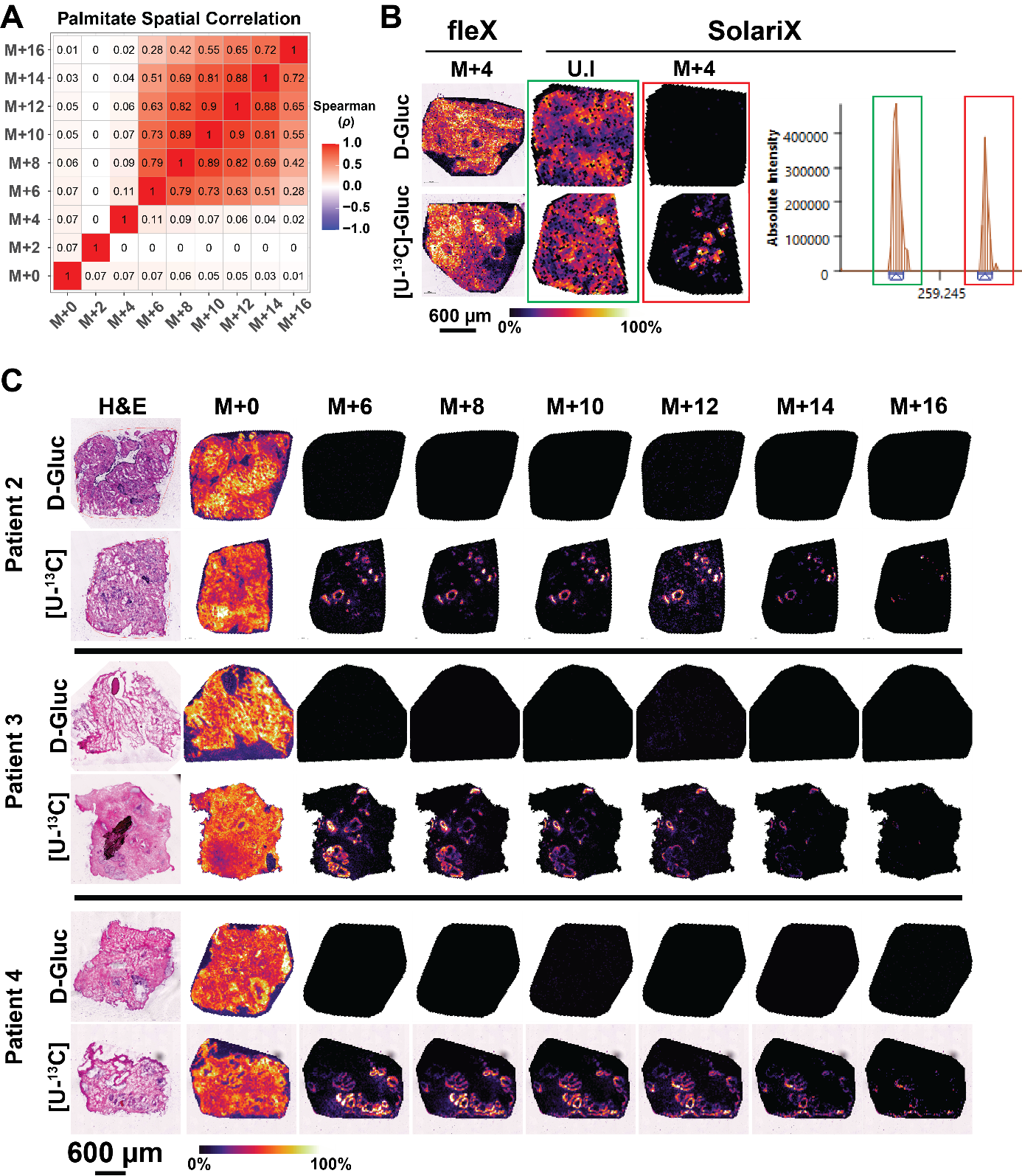
**Supplementary Fig. 1**

**Supplementary Fig. 1.** FA 16:0 isotopologue images are spatially correlated and overlap with endogenous analaytes can be resolved by high-resolution MS. A) Correlation heatmap showing the spearman correlation coefficient (p) for pairwise comparisons of FA 16:0 isotopologues in a representative tissue. B) Ion distribution images comparing the overlap of FA 16:0 M+4 between the timsTOF fleX and solariX FT-ICT platforms. Mass spectrum showing the baseline resolution of the M+4 signal and the endogneous unlabelled species. C) H&E and ion distribution images of FA 16:0 isotopologues from the remaining three patients in the cohort.

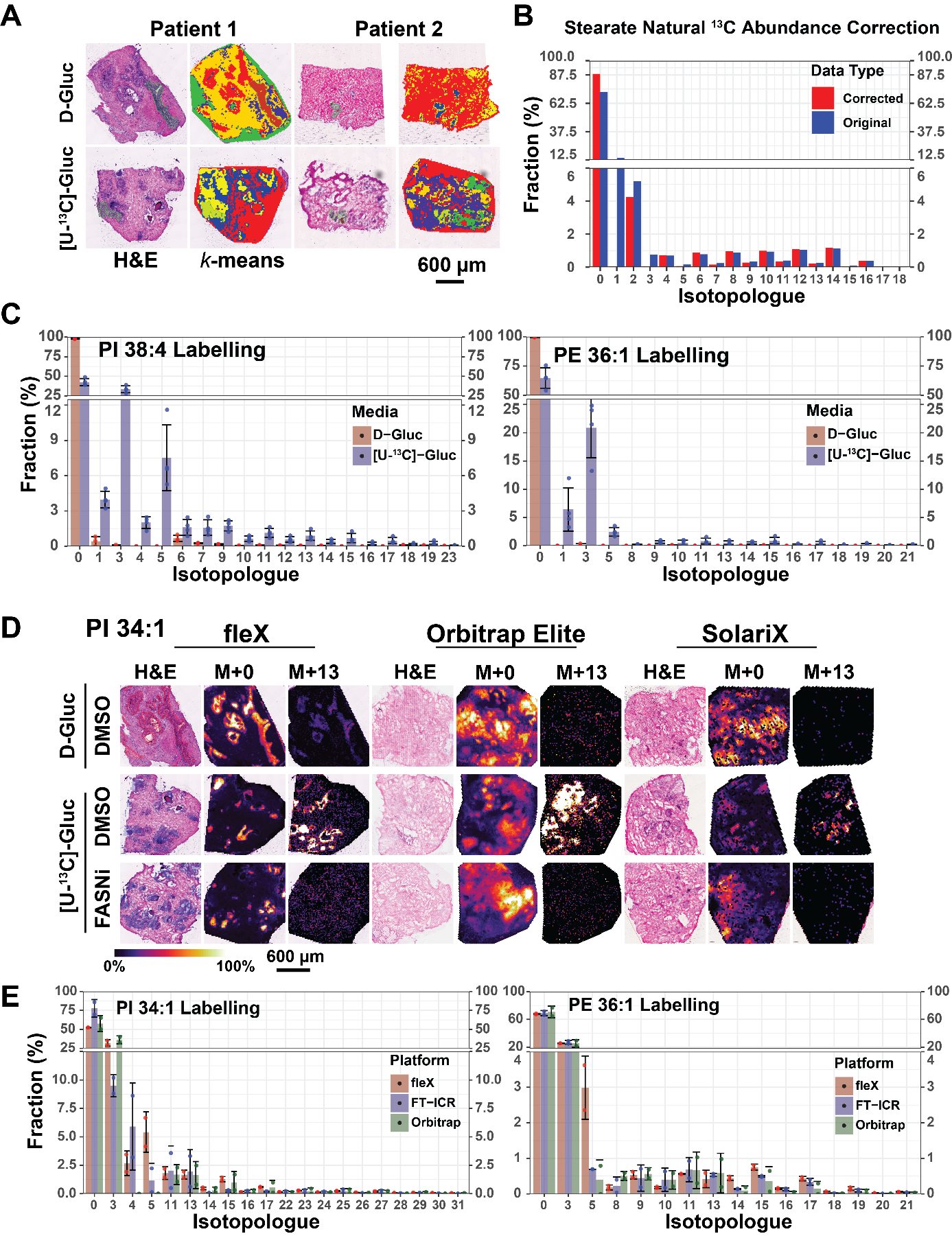
**Supplementary Fig. 2**

**Supplementary Fig. 2.** A) H&E stained images and bisecting k-means segmentation maps of two representative tissues cultured in D-glucose (top) or [U-^13^C]-glucose supplemented media. B) Labelling pattern of FA 18:0 before and after natural ^13^C isotope correction using IsoCor V2.2.3. C) Labelling patterns for PI 38:4 and PE 36:1 from PDE epithelia regions. Data represents average enrichment (%) ± SD of N=4 patients. D) Ion distribution images of PI 34:1 M+0 and M+13 with post-acquisition H&E stain from sections of the same patient tissue imaged across three MS platforms and labs. E) Isotopologue enrichment plot comparing measured fractions of PI 34:1 and PE 36:1 in a representative tissue treated with DMSO and cultured with [U-^13^C]-glucose across datasets acquired on different instruments. Enrichments were calculated based on isotopologues detected without across all three platforms.

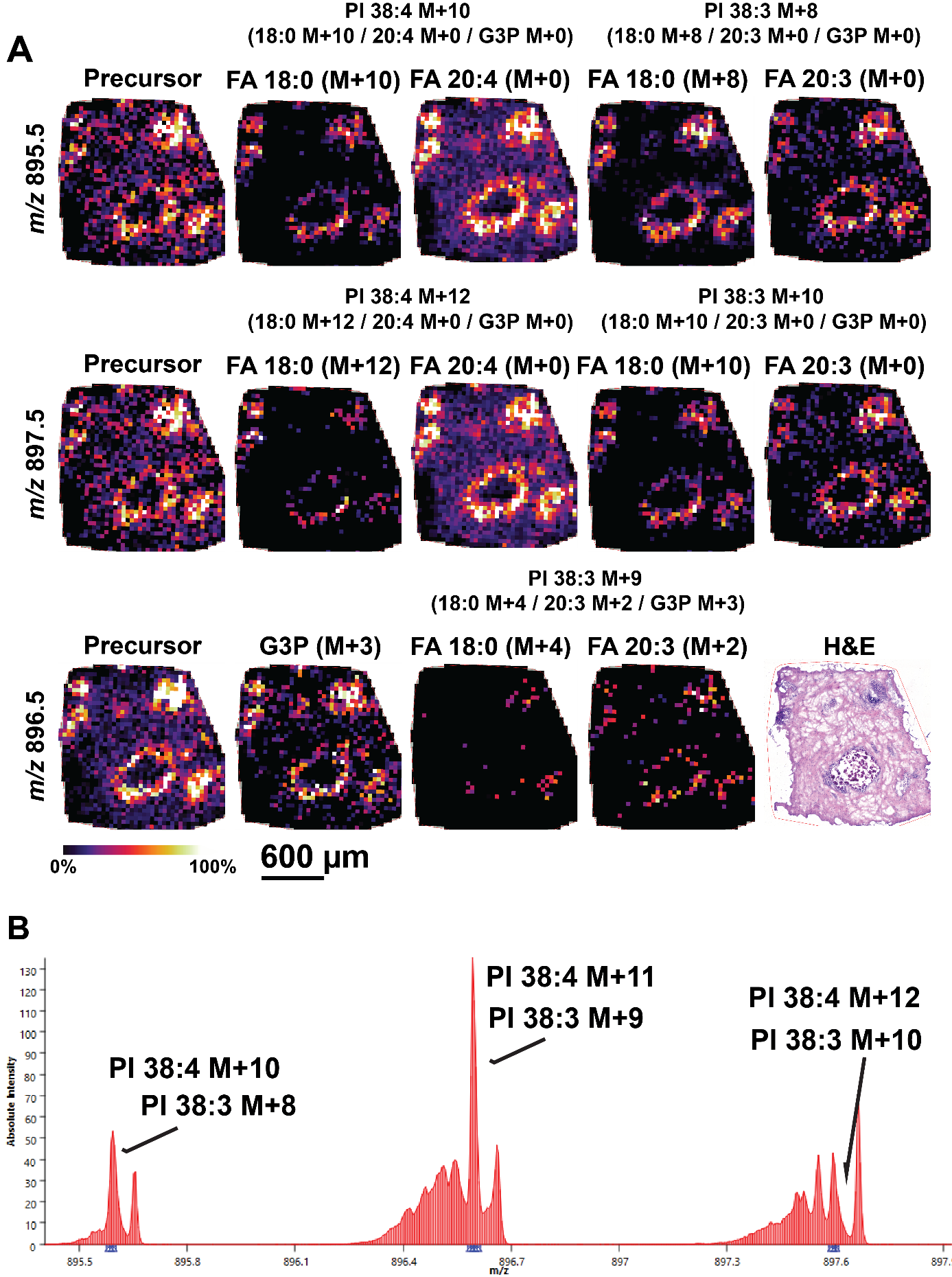
**Supplementary Fig. 3**

**Supplementary Fig. 3**. Targeted MS/MS imaging identifies distinct labelled PLs with varying degrees of FA labelling in prostate PDEs. A) Ion distribution images of precursors and labelled FA fragments annotating five distinct labelled lipids. B) Mean mass spectrum from MS/MS imaging data showing additional labelled lipids isolated in the quadrupole. All MS/MS imaging was performed at 40 µm pixel and step sizes. Ion distribution images are normalised within a single *m/z* channel.

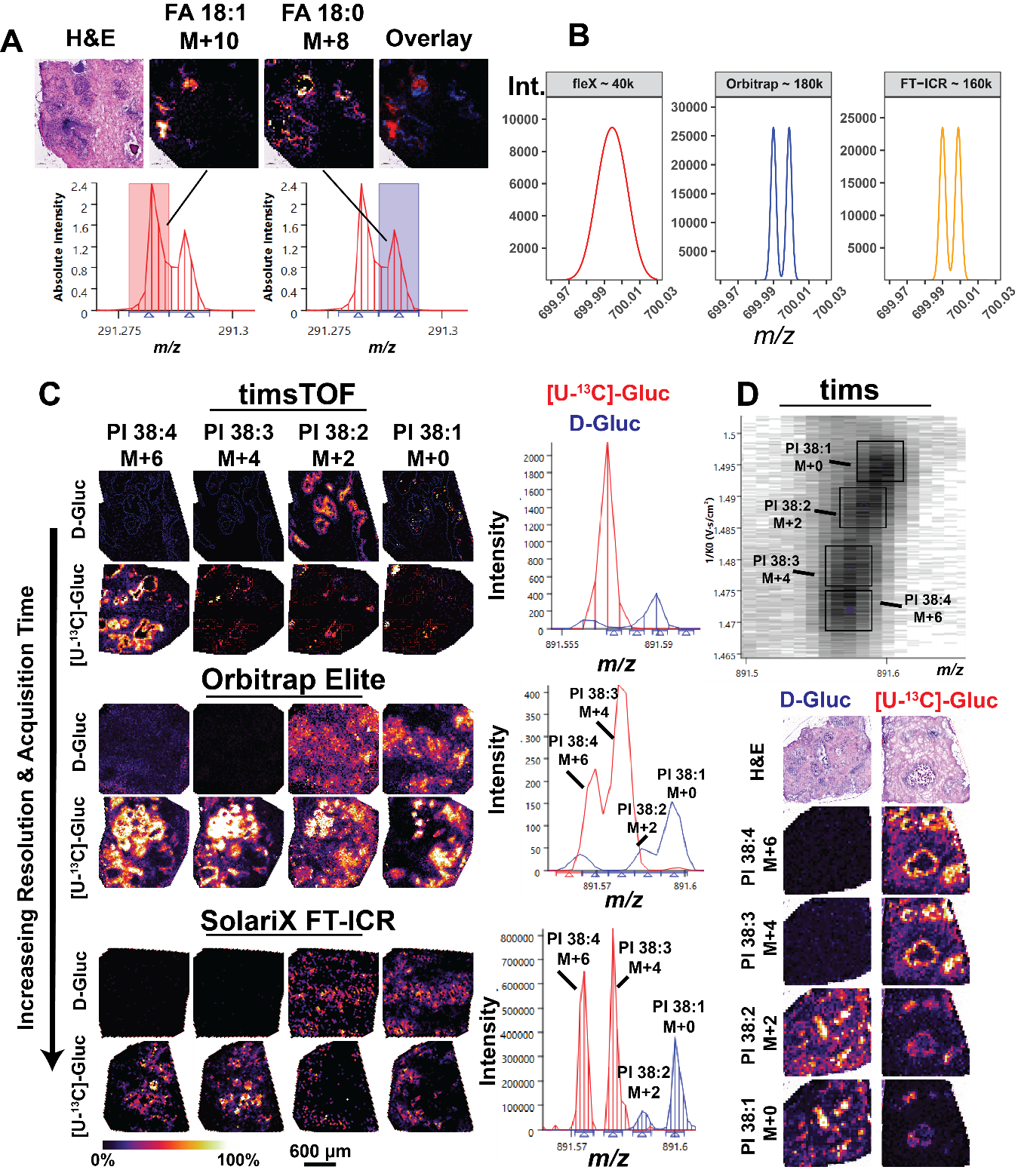
**Supplementary Fig. 4**

**Supplementary Fig. 4**. High-mass resolution MS platforms or orthogonal separation techniques resolve isobaric ^13^C-labelled lipids in prostate PDEs. A) Ion distribution images and mean mass spectrum of FA 18:1 M+10 and FA 18:0 M+8 in PDEs imaged on a timsTOF fleX. B) Theoretical resolution modelling showing the resolution of two theoretical labelled lipids differing my one double bond and two ^13^C labels at *m/z* 700.00 at resolutions expected across the three MS platforms used. C) Ion distribution images and mean mass spectrum of isobaric cluster at nominal mass *m/z* 891 containing four labelled PI lipids measured across three different MS platforms. D) Ion mobility heatmap and extracted ion mobility (EIM) images of the four labelled PI lipids. Ion distribution images are normalised within a single *m/z* channel.

**Supplementary Fig. 5**

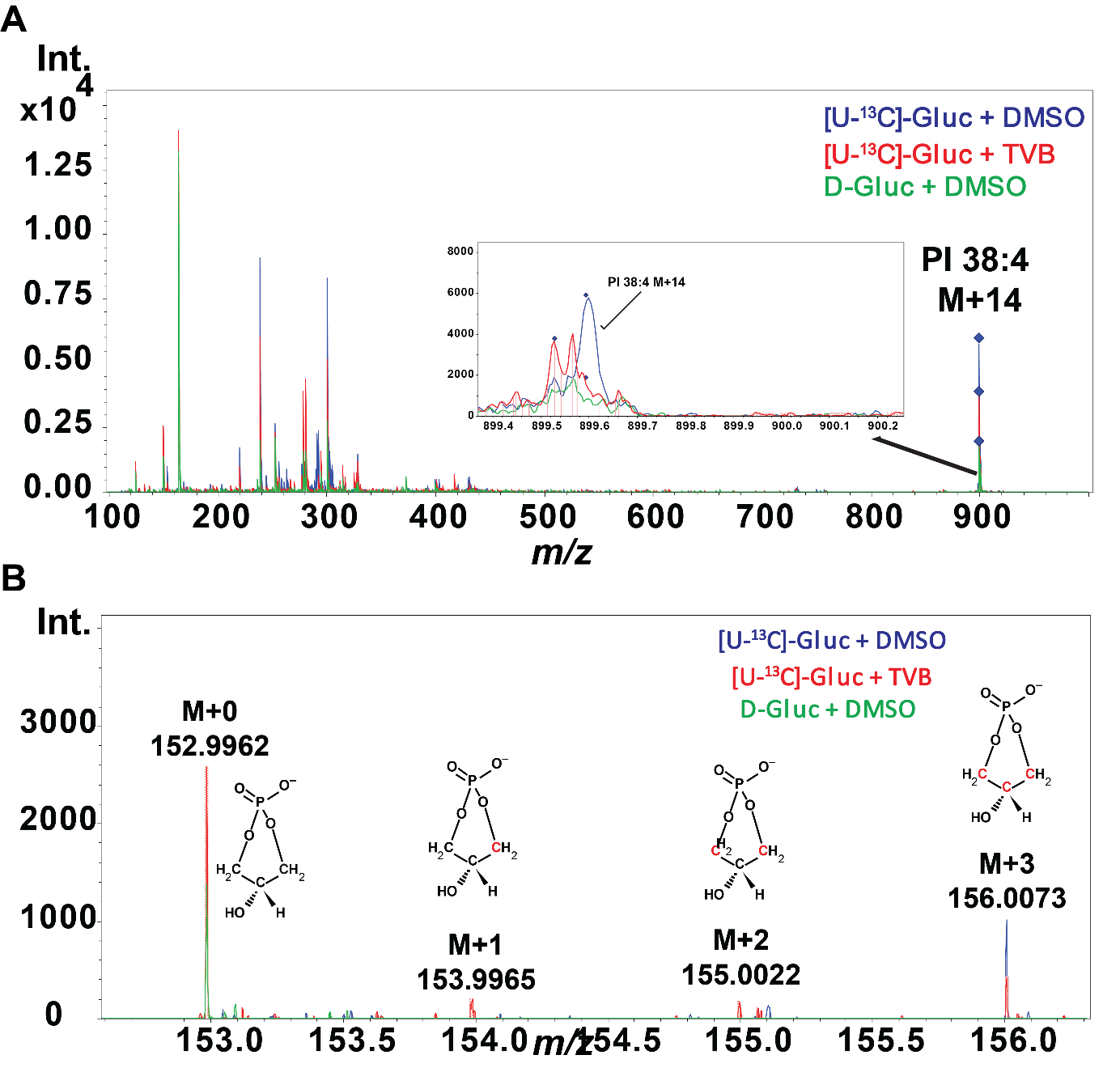

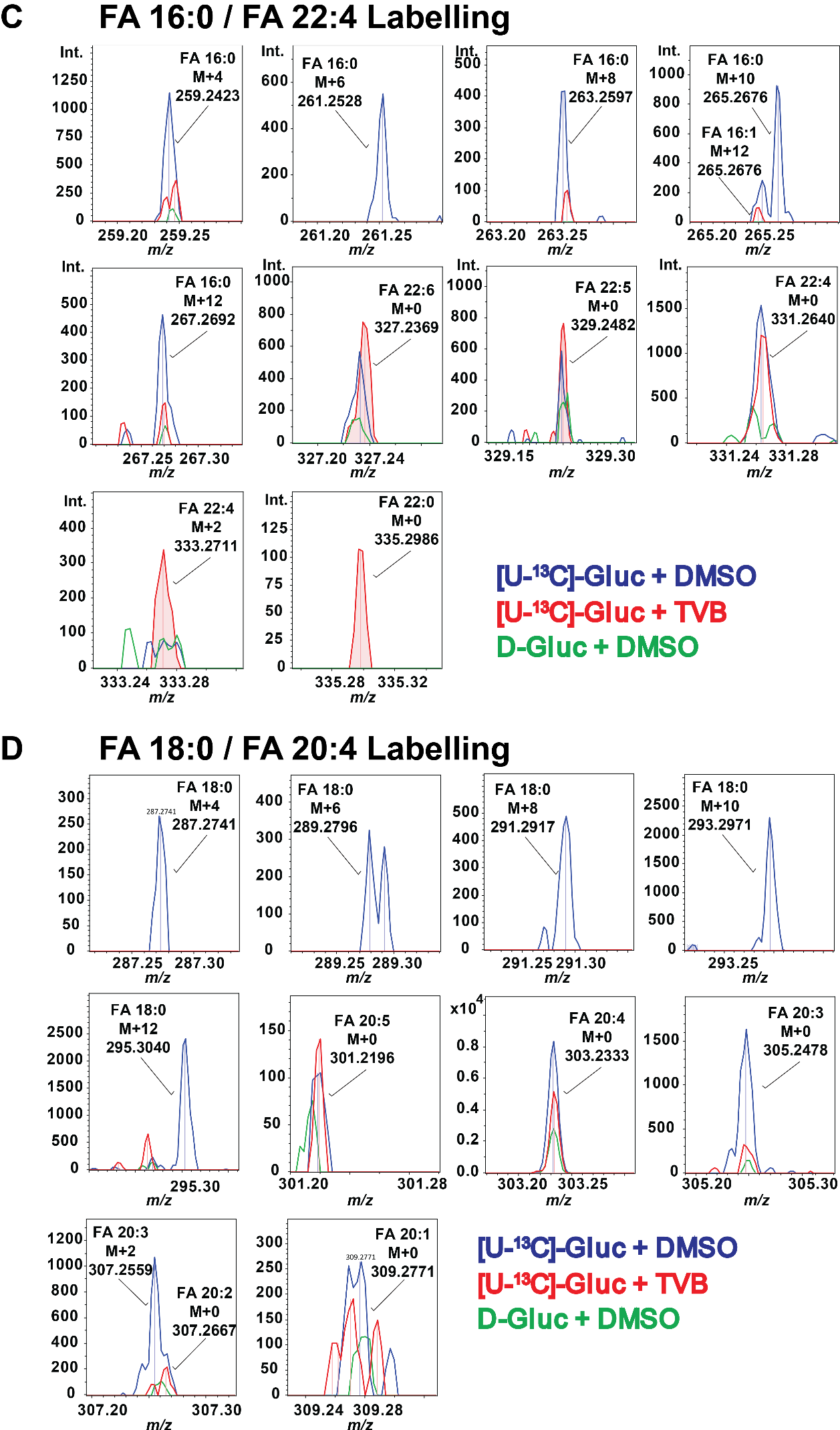

**Supplementary Fig. 5.** On-tissue MS/MS profiling reveals labelling of the G3P backbone in ^13^C-labelled lipids. A) Overlayed spectra of entire mass range acquired for MS/MS of precursor ion *m/z* 899.5968 representing the M+14 isotopologue of PI 38:4. B) Zoomed in region of the low *m/z* range (< *m/z* 200) showing fragment ions corresponding to G3P labelling.

**Supplementary Fig. 6**
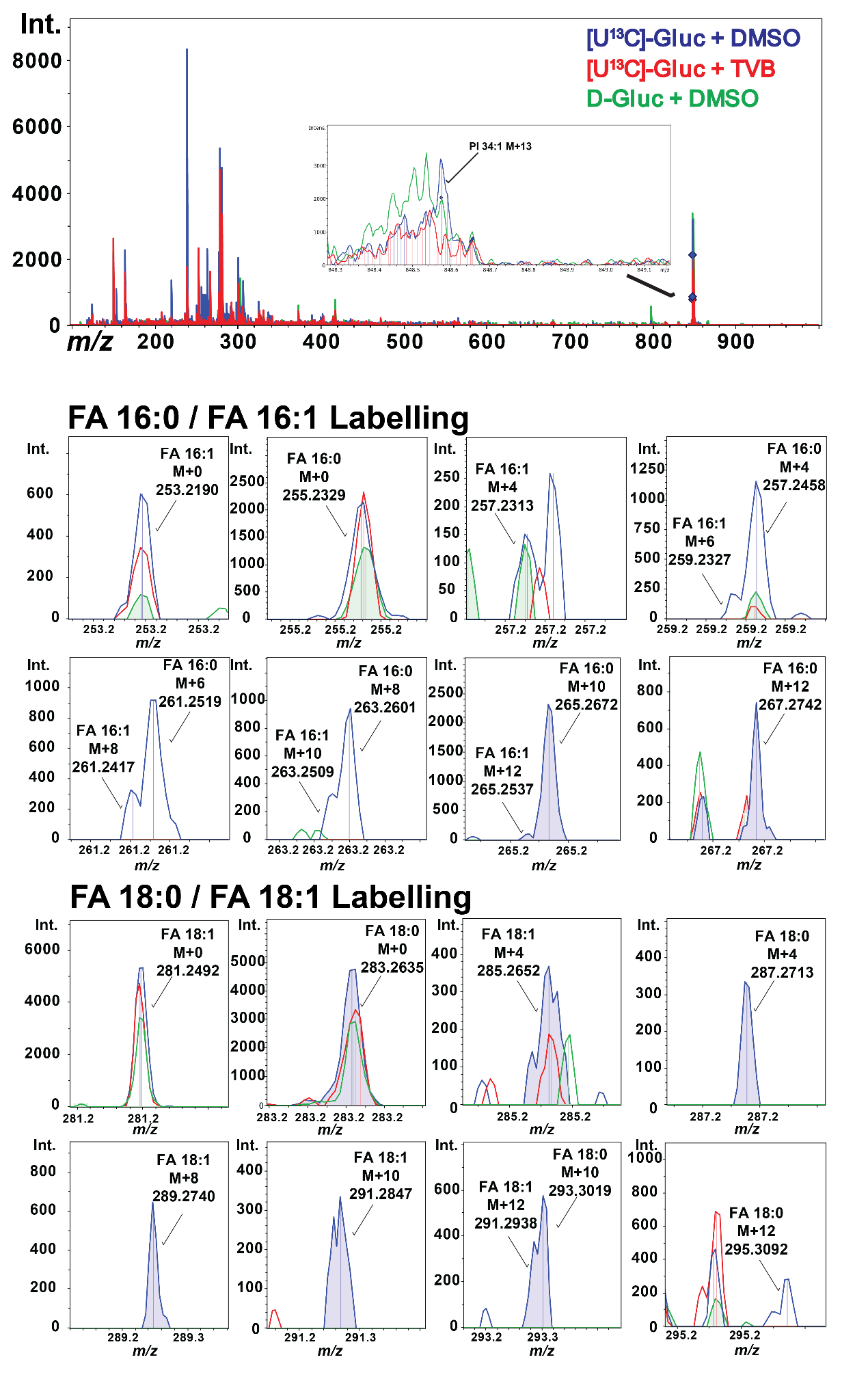

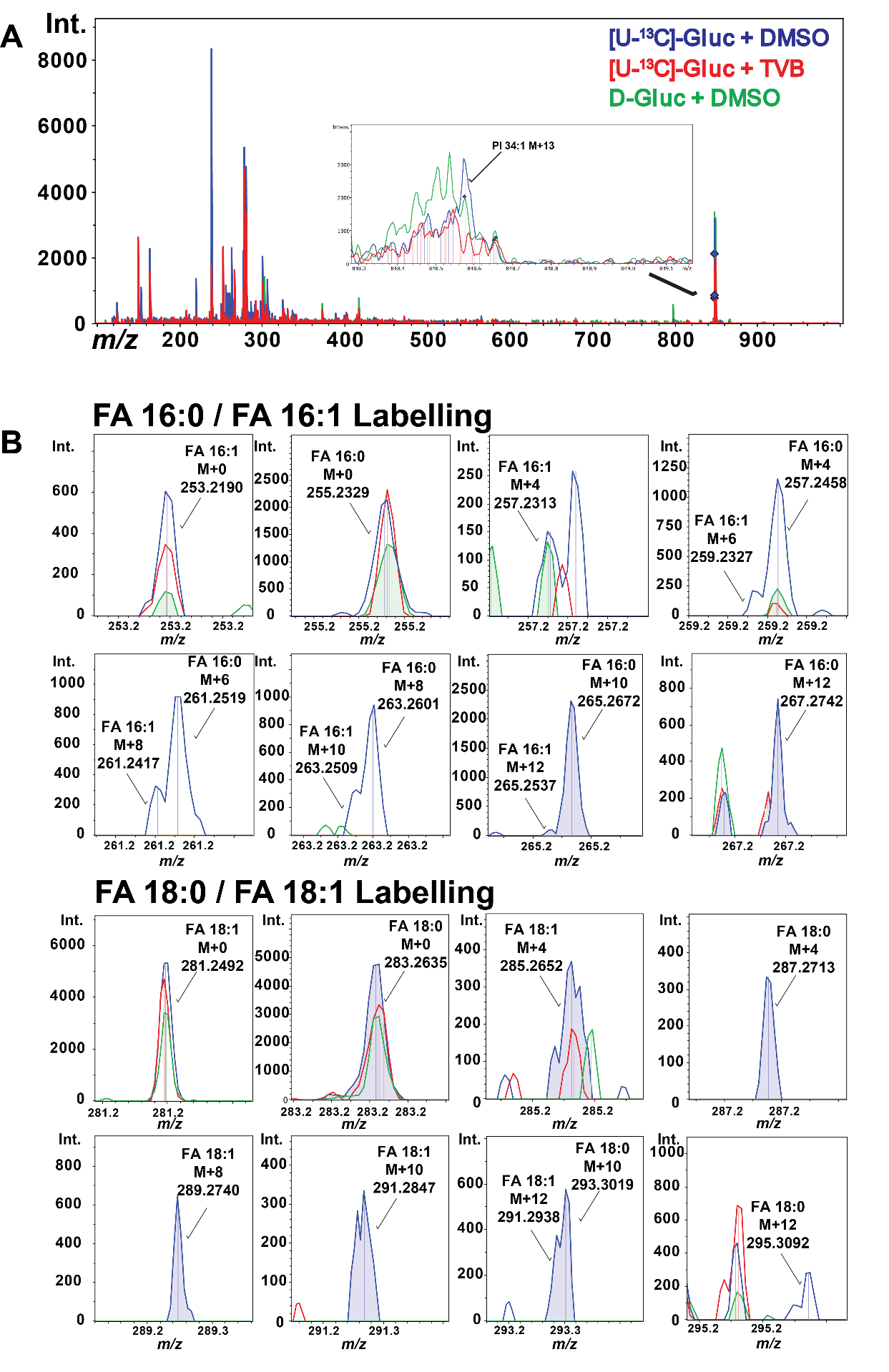

**Supplementary Fig. 6.** On-tissue MS/MS reveals heterogenous FA labelling patterns of ^13^C-labelled PI 34:1. A) Overlayed mass spectra from on-tissue MS/MS data of precursor ion *m/z* 848.5778 annotated as PI 34:1 M+13. B) Zoomed mass spectra showing labelling of different isotopologues of FA 16:0, FA 16:1, FA 18:0 and FA 18:1.

**Supplementary Figure 7**

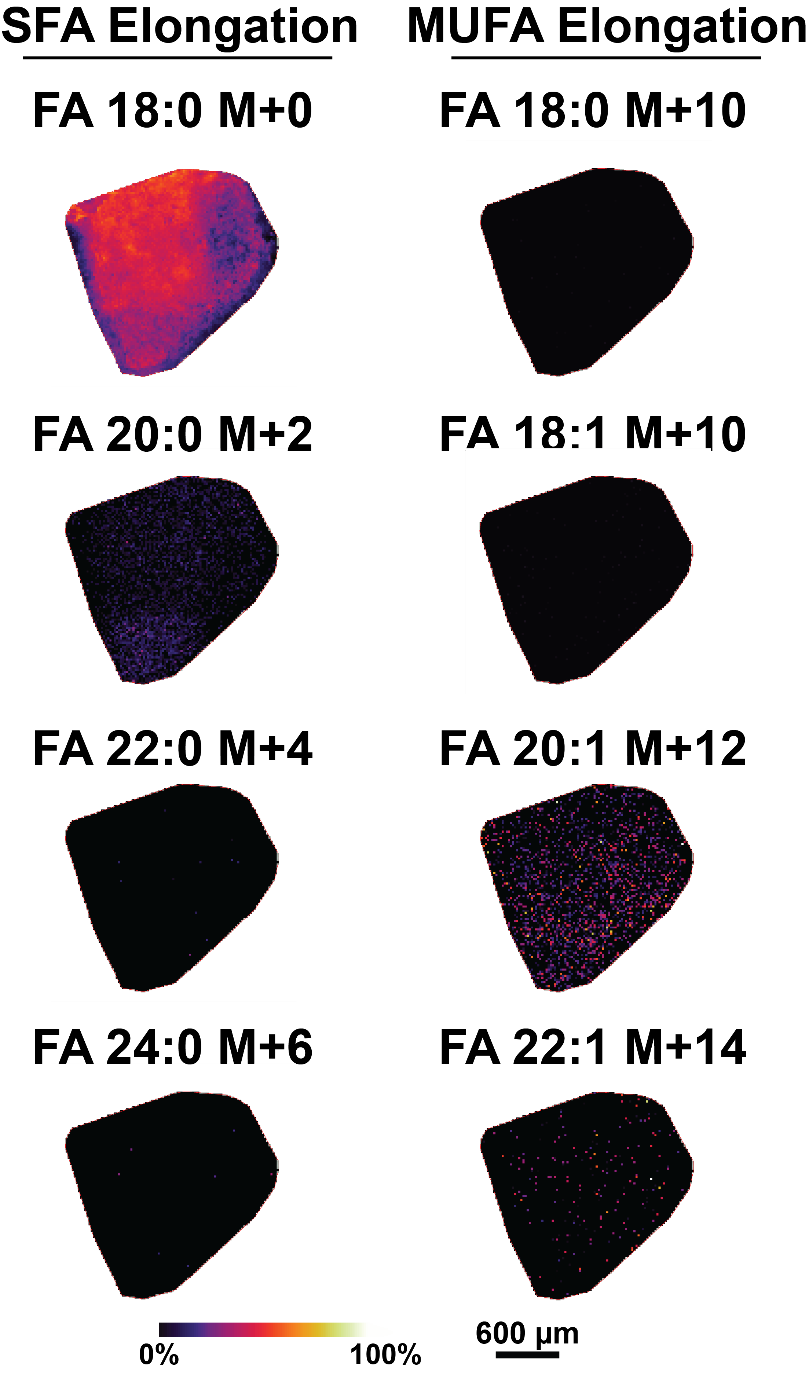

**Supplementary Fig. 7.** AIF MSI on matched control tissues show no interfering endogenous species generated through fragmentation. Ion distribution images of *m/z* values corresponding to labelled C20–C24 SFA and C20–C24 MUFAs measured in PDEs cultured with D-glucose supplemented media. Ion distribution images are normalised within each *m/z* channel.

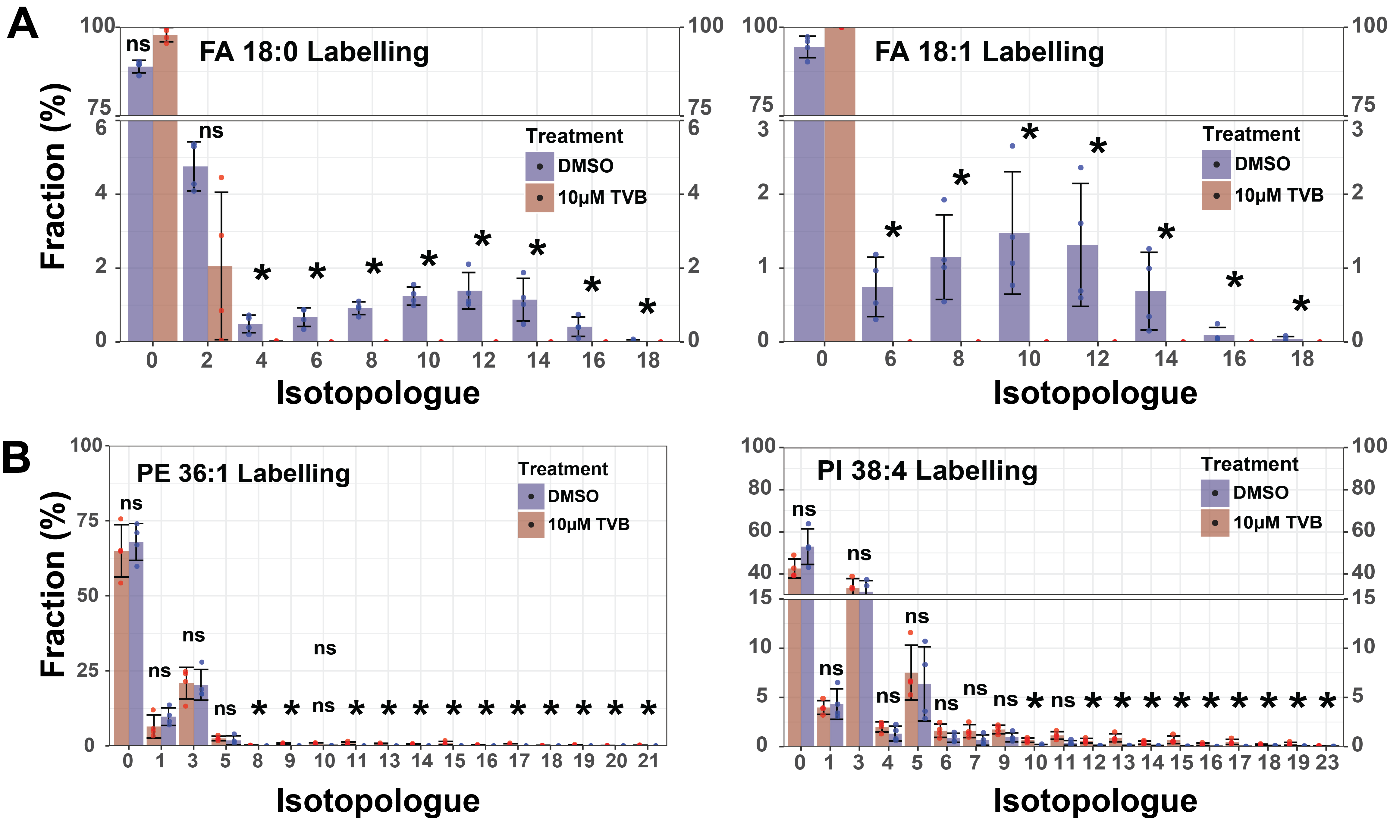
**Supplementary Figure 8**

**Supplementary Fig. 8.** PDE tissues treated with 10 µM TVB-2640 for 48 hours show a significant reduction in ^13^C-labelling for FASN-dependent labelling of A) de novo synthesised FAs and B) PI and PE lipids but no effect for FASN-independent labelling of G3P backbone. Data represents average enrichment (%) ± SD of N=4 patients and significance from Wilcoxon Rank Sum test with *p* <0.05.
